# Assessing and mitigating privacy risk of sparse, noisy genotypes by local alignment to haplotype databases

**DOI:** 10.1101/2021.07.18.452853

**Authors:** Prashant S. Emani, Gamze Gürsoy, Andrew Miranker, Mark B. Gerstein

## Abstract

Single nucleotide polymorphisms (SNPs) from omics data carry a high risk of reidentification for individuals and their relatives. While the ability of thousands of SNPs (especially rare ones) to identify individuals has been repeatedly demonstrated, the ready availability of small sets of noisy genotypes – such as from environmental DNA samples or functional genomics data – motivated us to quantify their informativeness. Here, we present a computational tool suite, PLIGHT (“Privacy Leakage by Inference across Genotypic HMM Trajectories”), that employs population-genetics-based Hidden Markov Models of recombination and mutation to find piecewise alignment of small, noisy query SNP sets to a reference haplotype database. We explore cases where query individuals are either known to be in a database, or not, and consider a variety of queries, including simulated genotype “mosaics” (composites from 2 source individuals) and genotypes from swabs of coffee cups from a known individual. Using PLIGHT on a database with ~5,000 haplotypes, we find for common, noise-free SNPs that only ten are sufficient to identify individuals, ~20 can identify both components in two-individual simulated mosaics, and 20-30 can identify first-order relatives (parents, children, and siblings). Using noisy coffee-cup-derived SNPs, PLIGHT identifies an individual (within the database) using ~30 SNPs. Moreover, even when the individual is not in the database, local genotype matches allow for some phenotypic information leakage based on coarse-grained GWAS SNP imputation and polygenic risk scores. Overall, PLIGHT maximizes the identifying information content of sparse SNP sets through exact or partial matches to databases. Finally, by quantifying such privacy attacks, PLIGHT helps determine the value of selectively sanitizing released SNPs without explicit assumptions about underlying population membership or allele frequencies. To make this practical, we provide a sanitization tool to remove the most identifying SNPs from a query set.

## Introduction

Privacy concerns in the digital age are ubiquitous, with individual data collection, access, and the sophistication of tools of attack constantly increasing to render individuals vulnerable to a significant risk of compromising data exposure. Incursions upon individual privacy include the removal of personal control over the uses of such data, and the possibility of being subjected to discrimination on the basis of revealed information. Perhaps the most invasive forms of such attacks involve gaining access to information on the physical and mental constitution of an individual without their knowledge or consent. Such breaches are becoming increasingly likely in an era marked by massive health-based data collection and digitization efforts, whose ultimate goals include the provision of personalized medical interventions. Genetic data lie at the heart of these tailored medical approaches, as many phenotypes are believed to have an ultimate basis in our genetic makeup, and as such, are being collected as a part of large-scale projects such as the UK Biobank (www.ukbiobank.ac.uk) and the NIH’s AllofUs (allofus.nih.gov) program.

Based on essential work by Homer et al (Homer et al. 2008), genetic data in general, and single nucleotide polymorphisms (SNPs) in particular, were shown to enable the identification of individuals in DNA mixtures. Additional work reaffirmed the ability of SNPs to reveal whether an individual belonged to a study cohort or DNA mixture (Visscher and Hill 2009). Gymrek et al (Gymrek et al. 2013) exploited the possibility of linking the genomes of individuals to surname data from genealogy websites, thereby clearly demonstrating the risk of exposure from the public release of genotype data. Nowadays, law enforcement agencies frequently employ SNP data in the identification of individuals, and their genetic relatives. While it has been suggested 20-30 independent SNPs are enough to re-identify individuals (Lin et al. 2004), this quantification needs to be updated based on available databases of human genomes and auxiliary biological data.

The increasing availability of cheap sources of SNP extraction are a cause for concern. SNPs can be extracted from genotyping and omics assays, say, as a part of genetic studies of disease vulnerability; forensics analyses of found objects; and can be inferred from established genotype-phenotype relationships. Functional genomics assays in particular allow both direct sequence-based extraction of variants (Gürsoy et al. 2020), as well as indirect inference based on gene expression values and associated loci (Harmanci and Gerstein 2016), and the ubiquity of large-scale omics projects make this source of variants especially concerning (Gürsoy et al. 2020). Several studies have demonstrated the risk of identification even in the case of partial privacy preservation measures such as SNP beacons (Shringarpure and Bustamante 2015; Raisaro et al. 2017; Von Thenen et al. 2019), and the publication of only GWAS summary statistics (Im et al. 2012). For some time, there has been a strong case for the restriction of access to genotypic data and comprehensive privacy preservation, say, through encryption-based analysis (Erlich and Narayanan 2014); however, in the case of genotypes that can be inferred from functional genomics data or environmental samples, the field is now directing attention towards the notions of data sanitization (Gürsoy et al. 2020).

All arguments in favor of protectionism are frequently confronted by the scientific community’s desire for unrestricted access to datasets: undoubtedly, increased public access to data would democratize information, and enable biological analyses of greater statistical power in proportion to the size and quality of the available data. Striking a balance between the two requires a clear quantification of the risks of re-identification and inference, relative to the proposed benefit of releasing the data. In response to this challenge, we provide a computational tool that assesses the degree to which a set of released SNPs could lead to genotypic and phenotypic inferences, using a Hidden Markov Model (HMM) approach (**Figure 1**). The tool is termed “**P**rivacy **L**eakage by **I**nference across **G**enotypic **H**MM **T**rajectories” or *PLIGHT*.

**Figure 1.**
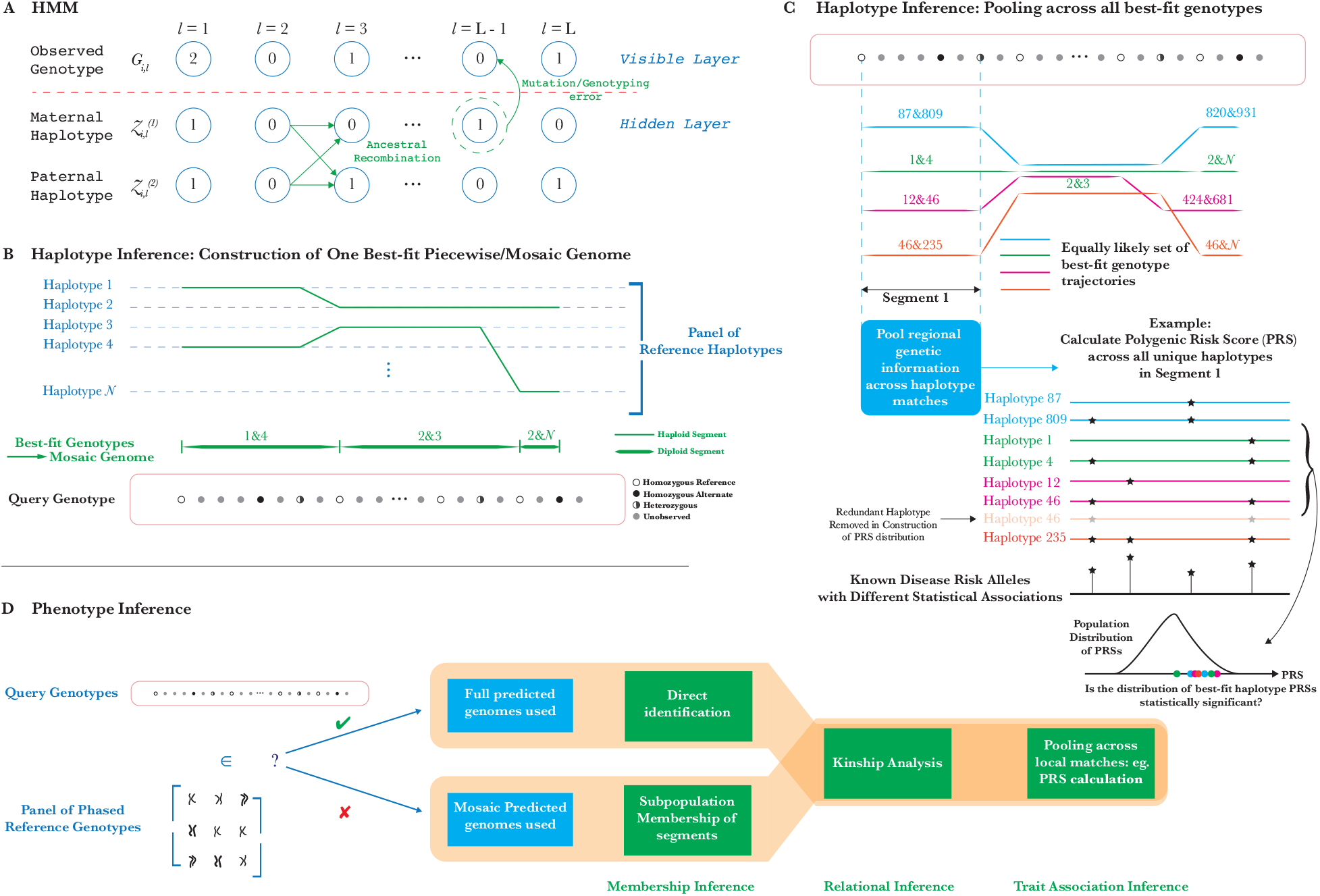
Methodology of the HMM-based and inference analyses in PLIGHT. (A) Schematic of the Li-Stephens HMM model for haplotypic recombination and mutation. (B) Inference of best-fit haplotype pairs in different genomic regions based on a chosen database of reference haplotypes, and the subsequent construction of the diploid mosaic genome. (C) An example of how the haplotype inference procedure described here could be used for novel attacks. By calculating polygenic risk scores (PRSs) for particular traits across all identified mosaic genome segments, we can assess whether there is a statistically significant clustering of the PRSs relative to background values. If this is found to be the case, the individual is more likely to be linked to such a trait. (D) Exploration of the types of phenotypic inference attacks possible, conditional on whether the individual is known to be in the database of reference haplotypes or not.

The premise of the tool is that even limited, noisy and sparsely distributed genotypic information carries with it a certain risk of identification and downstream inference. Concerns about noisy and sparse SNPs, in particular, are legitimate due to the ease with which they can be obtained. If a set of noisy SNPs could be assessed indirectly through inference, or through direct access to genetic material acquired from objects in contact with an individual (Gürsoy et al. 2020), we show that it is often possible to expand upon this partial information using existing genetic databases. For example, consider the case where a single DNA fragment can be separated from contaminated genetic material, thus reliably ensuring a single source individual. With the advent of long-read technologies such as nanopore (Jain et al. 2016) and SMRT (Eid et al. 2009) sequencing, we can now read that fragment to a length of hundreds or thousands of base pairs. The data is somewhat error-prone due to the dearth of material, but still could yield a few noisy SNPs along the length of the fragment. Additionally, certain physical characteristics, and knowledge of Mendelian disorders in individuals, potentially leak information on mutations at particular loci (Posey et al. 2019). The assessment of identification risk for limited SNP data is especially important in light of the availability of large-scale public genotypic databases of individuals stratified based on geographical and putative ethnic groupings, as well as SNP sets associated with (potentially socially compromising) disease risk.

Inspired by imputation methods such as IMPUTE2 (Howie et al. 2009) and Eagle (Loh et al. 2016), the inference procedure in PLIGHT is based on the Li-Stephens model (Li and Stephens 2003), where an HMM is used to explore the space of underlying pairs of haplotypes in a diploid genome with the possibility of *de novo* mutations and recombination between haplotypes. In contrast to imputation, however, PLIGHT does not identify the most likely SNPs at each genomic locus (which would be an underconstrained problem for very sparse SNP sets). Rather, a solution to the inference problem consists of finding which segments of existing haplotypes in a database best match the query SNP set. These segments are pieced together to form a continuous path, or genotypic *trajectory*, through the reference haplotype space matching the full query SNP set. Thus, while phasing and imputation already have highly efficient solutions to find the single most likely genotype at an unobserved locus based on sufficiently dense nearby SNPs, our method is optimized to find all equally likely trajectories pieced together from reference haplotypes.

Several studies in the field of genetic privacy have considered k^th^-order correlations between SNPs (k=2 corresponds to pairwise linkage disequilibrium (LD)) to maximize the probability of reidentifying individuals (Von Thenen et al. 2019). PLIGHT seeks to capture all these higher order correlations, using the pre-existing correlation structure (of *any* order) of the reference genetic database with a biologically plausible model of recombination (the default value being based on HapMap Consortium data (Li and Stephens 2003; Marchini and Howie 2010; International HapMap Consortium 2003), though users can add their own custom, position-specific recombination rates. PLIGHT is agnostic to any assumptions of membership of the query individual to a specific subpopulation, population homogeneity or to estimates of allele frequencies; rather, it is dependent on the composition of the whole reference database. Additionally, we leverage this inference procedure to make a more informed choice of which SNPs to sanitize, in the case of inferred genotypes from data whose primary purpose is not the identification of variants (e.g. RNA-seq or ChIP-seq data). We include the sanitization tool in the PLIGHT suite.

The employment of HMMs in the modelling of LD in genomes has substantial precedent, such as in the inference of local ancestry (Pasaniuc et al. 2009; Price et al. 2009; Baran et al. 2012) and the determination of regions that are Identical by Descent (IBD) (Bercovici et al. 2010). For example, Baran et al (Baran et al. 2012) use a latent-state-space reduction by running a two-level HMM: an inner model for each ancestral population within a genomic window of a certain length and a higher-level model for exchanges between the windows. We choose a more straightforward implementation of HMMs, to avoid any consequent assumptions either of the density of query SNPs (defining a genomic windows precludes recombination between SNPs in that window) or of ancestral membership. There is, however, a consequent price paid in terms of an increase in running time and memory usage. We partially ameliorate these costs using sampling and pooling approaches described below. We also note that the Positional Burrows-Wheeler Transform (PBWT) (Durbin 2014) has been used in tandem with the Li-Stephens model to improve the efficiency of haplotype matches across large databases (*fastLS* (Lunter 2019)). The scaling of this method with database size is far superior to the straightforward HMM implementation. However, for now, *fastLS* does not allow position-specific variations in the recombination rate. For sparsely distributed SNPs, there will be large variability in the recombination rates between any pair of adjacent observation sites. PLIGHT allows for variation of the recombination model, and even include position-dependent recombination effects. Further, in contrast to the PBWT formalism, PLIGHT also enables the user to include trajectories that are less-than-optimal, for robustness checks or exploration of noisy data.

In summary, the primary aims of this study are: (1) To demonstrate that even a few SNPs leak considerable information about source individuals, not just by increasing the capacity to find them or genetic relatives in databases, but also by allowing matches of subsegments of their genotypes to reference databases; (2) To provide tools to quantify these varying degrees of information leakage and to subsequently sanitize query SNPs in a manner that balances utility of datasets and privacy of the source individuals.

## Results

### Repurposing population-genetics models for privacy

The primary purpose of PLIGHT is to serve as a tool to quantify SNP information content, and leverage that information to make judicious choices on SNP masking (*sanitization*) from datasets. The expectation is that an attacker would acquire a set of unphased alternate allele dosages {0,1,2} at particular SNP loci. Inference by means of a reference genetic database requires that the SNP position overlap, at least partially, those that have been genotyped in the database. Each identified SNP may have a unique probability associated with the alternate allele dosage (reflecting certainty) or an overall error rate may be assigned. The Li-Stephens HMM Model (**Figure 1A**) then maps the recombination and mutation processes onto the transition and emission probabilities, respectively. In this way, the process of aligning a set of query genotypes can be seen as simultaneously phasing the genotype into the paternal and maternal haplotypes at a single query locus, and allowing for transitions to other haplotypes between adjacent query loci. In doing so, segments of the query SNPs are found to match to pairs of haplotypes which may transition to other pairs for the next segment (**Figure 1B**).

Formally, we find the trajectories as sequences of reference haplotype pairs (for a diploid genome) at each locus that best fit the observations: that is, for observed query SNPs at genomic loci *l* = {1,2, …, *L*}, the trajectory 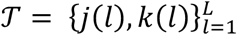, where *j* and *k* are the labels of best-fit reference haplotypes as a function of the observed genomic position (**Supplemental Fig S1**). We label a single best-fit trajectory as a diploid *mosaic* genome if the pairs of matching reference haplotypes change across the observed loci. The collection of all equally likely trajectories for a given query SNP set is the primary output of the algorithms.

Having access to all equally likely trajectories may enable novel forms of attack (**Figure 1C**), which we explore in subsequent sections. PLIGHT also provides a visualization of all trajectories across the observed loci, and the logarithms of the joint probabilities of observing the query SNPs for: (a) the HMM model, (b) the case where all SNPs are independent and satisfy Hardy-Weinberg equilibrium (described by their MAFs) and (c) the case where all SNPs are independent, but are described by their genotype frequencies (and not their MAFs). The log probability of the most likely trajectories are also presented. Data publishers then have the option of removing (“sanitizing”) the most informative SNP from the observed sample, based on identifying segments with the smallest diversity of inferred trajectories (i.e. segments with the fewest trajectories passing through them) and the SNP allele frequencies.

A major consideration behind the design of our methods was contending with the scale of reference genotype databases: with the search space for matches to query genotypes ranging from thousands to (eventually) tens or hundreds of thousands of genotypes, HMMs quickly become computationally intractable in terms of the required memory and speed. Our approach to the problem was to construct three algorithms that negotiate the tradeoff between exactness of the calculation and the burden on memory resources:

1. *PLIGHT_Exact* performs the exact HMM inference process using the Viterbi algorithm (Viterbi 1967).
2. *PLIGHT_Truncated* was inspired by the state-space reduction methods in the Eagle2 imputation program (Loh et al. 2016), and phases in a process of truncating the set of all calculated trajectories to only those within a certain probability distance from the maximally optimal ones, resulting in a smaller memory footprint.
3. *PLIGHT_Iterative* iteratively partitions the reference search space into more manageable blocks of haplotypes and runs *PLIGHT_Exact* on each block, followed by pooling and repetition of the scheme on the resulting, smaller cohort of haplotypes. This algorithm has significantly better scaling properties than the others, with respect to the size of the reference database.

Thus, *PLIGHT_Truncated* and *PLIGHT_Iterative* are approximations to the full state space exploration designed to reduce hard disk memory usage, and both hard disk and RAM usage, respectively. However, for most purposes *PLIGHT_Truncated* is superseded by *PLIGHT_Iterative* in performance. *PLIGHT_Truncated* mainly serves to determine the “compressibility” of the trajectories, i.e. the size of the trajectory subspace that is sufficient to match the results of the exact algorithm.

Subsequently, the results of all algorithms are visualized using a visualization module called *PLIGHT_Vis*, and processed using downstream inference modules. If the user then decides to remove a SNP on the basis of these results, they can employ *PLIGHT_SanitizeGenotypes* to do so: a SNP is eliminated from the query set, which is then rerun through one of the three HMM algorithms to produce the sanitized results. The overall computational framework is depicted in **Figure 2**.

**Figure 2.**
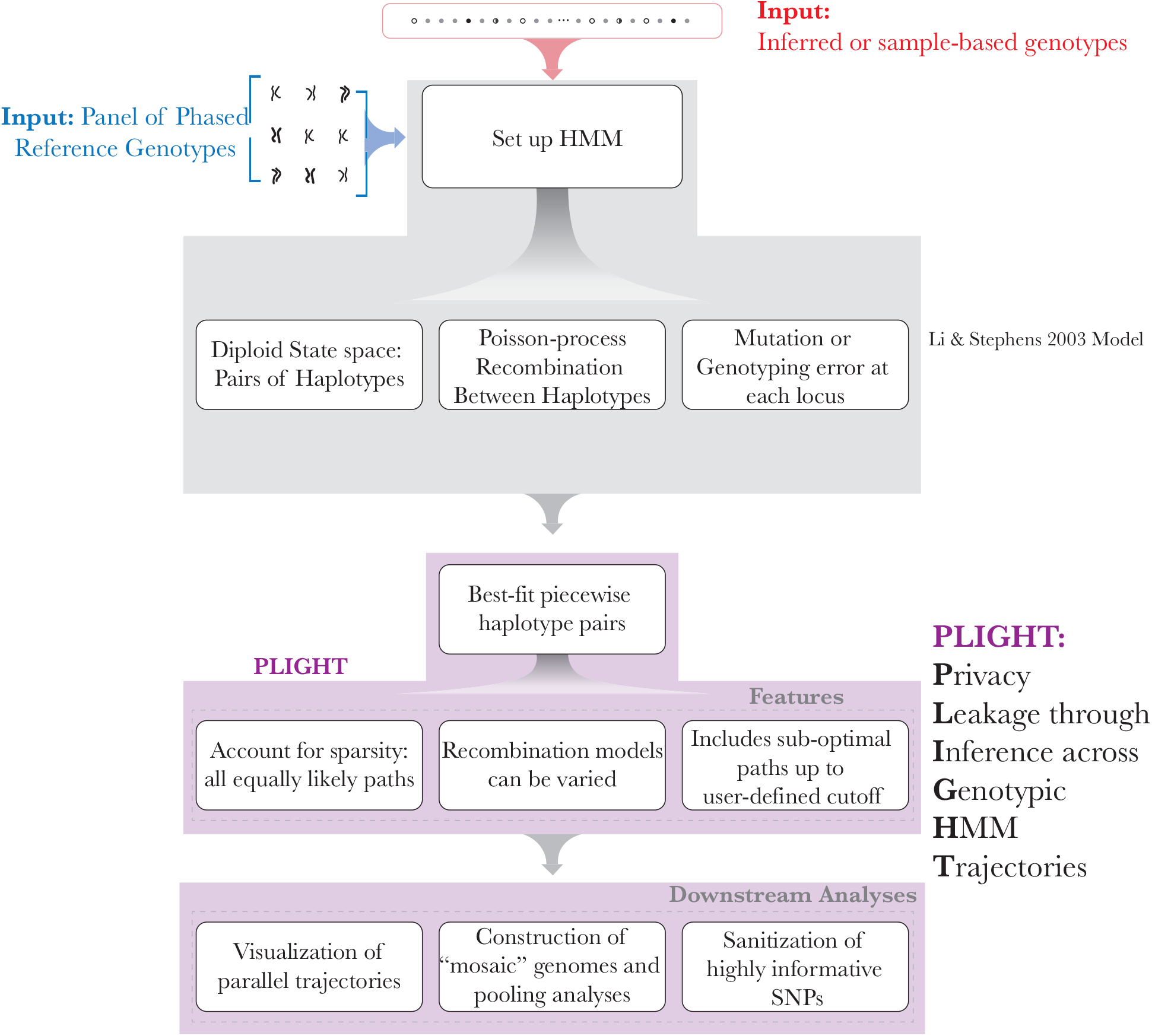
Computational framework of PLIGHT. The inputs of a reference database and of the query SNPs are shown at the top. Next, the population-genetics model by Li and Stephens forms the core of the HMM-based framework. This includes the definition of a diploid state space, a Poisson-process based recombination model with rate of growth proportional to the genomic distance between two loci, and a mutation/error rate at each locus. This model is implemented within PLIGHT, which identifies the best-fit reference haplotype labels in the diploid state space. Special features include the identification of all equally likely trajectories, flexibility in the recombination models used, and an allowance for sub-optimal trajectories to be identified. Furthermore, PLIGHT includes visualization and SNP sanitization modules. We show how the framework also allows for the construction of mosaic genomes and analyses involving the pooling of information across all identified mosaic trajectories.

### Attack Scenarios

In this section, we lay out a few examples of the real-world scenarios in which we reasonably expect to observe a small number of noisy SNPs, the types of attacks that may be carried out using those SNPs (**Figure 1D**), and the applicability of PLIGHT to those attack scenarios.

#### Sources of small numbers of SNPs

##### 1. Low quality and /or contaminated samples

DNA samples acquired from contacted objects (such as coffee cups) can in general yield thousands of SNPs (see for example, (Gürsoy et al. 2020)). However, if DNA samples become damaged or contaminated by the presence of other DNA sources, the yield of high-confidence SNP calls may be significantly reduced (Tillmar et al. 2018; Turchi et al. 2019; de Vries et al. 2022). It is important to explore the lower bounds of SNP numbers that would suffice for certain types of privacy attacks, given the ease and legality of surreptitiously extracting variants from environmental DNA samples.

##### 2. Nanopore reads

In cases of samples containing DNA from multiple individuals, nanopore sequencing technologies could be employed to read individual DNA strands. A single read from nanopore sequencing would reliably belong to a single individual (thus removing the effects of contamination), and may yield a handful of noisy SNPs.

##### 3. Inferred SNPs

Although more speculative, we consider the possibility of sourcing SNPs from knowledge of individual characteristics. If even a small number of SNPs can be guessed, say on the basis of known Mendelian disorders in an individual or their genetic relatives, it might be possible to piece together sufficient genetic information to find such individuals (or relatives) in sensitive phenotypic study databases. Inference of SNPs has also been demonstrated for derived genomic data, such as expression quantitative trait loci (eQTLs) (Harmanci and Gerstein 2016) and allele-specific expression (Gürsoy et al. 2021). Again, the number of inferred SNPs may be considerable, but with varying degrees of confidence. The highest quality matches would form a smaller subset. Finally, a small number of germline SNPs may be leaked in somatic variant call-sets (Sendorek et al. 2018; Meyerson et al. 2020), though evidence is ambiguous as to the number of true germline variants.

#### Types of privacy attacks considered

##### 1. Membership inference attack

The identification of an individual as a member of a database or cohort could allow the linking of that individual to potentially stigmatizing phenotypes. Moreover, we specifically consider two forms of membership inference:

a. *Full membership:* This is the case when an individual is found to be part of a database.
b. *Partial membership:* This is the case when partial segments of the query individual’s genome are found in a database. This could occur due to Identity-by-Descent (IBD) or Identity-by-State (IBS). Partial membership allows certain regions of the genome to be related to a phenotypic cohort; e.g. the query individual may overlap in certain genomic regions exclusively with individuals in the phenotype-positive group (i.e. with cases and not controls).

##### 2. Relational (or Kinship) inference attack

This type of attack technically overlaps with the partial membership attack mentioned above, but warrants its own discussion. Partial matches in certain genomic regions could allow the genetic relatives of a query individual to be found in phenotypic databases, thus exposing them to an attack on their privacy.

##### 3. Trait association inference attack

This type of attack involves using the partial information from regional matches to the genome to make inferences about the individual. Consider the following hypothetical example. An individual is in contact with an object. The attacker collects DNA from the object, identifies the subset of high-quality SNPs, and proceeds to infer aspects of the individual — such as which genetic subpopulation the individual belongs to, or whether the individual has GWAS SNPs or a polygenic risk score (PRS) that is high for a particular phenotype.

#### Deploying PLIGHT to address attacks

We use unphased SNPs with a user-defined level of confidence/error rate as inputs to PLIGHT tools. We demonstrate full membership attacks (a) in the presence of genotyping error and (b) when SNPs from other individuals contaminate a primary source. The full membership attack is the simplest form of HMM inference, because we restrict the recombination rate to be 0 (and so the transition probabilities are 0). This simple HMM model is still different from the assumption of independent SNPs because potential correlations exist between query SNPs across the genetic database.

We also search for relatives of query individuals, by combining partial matches to the genomes across chromosomes. Finally, we explore a version of the trait association attack where we use the matched segments (for individuals not in the database) to partially impute SNP dosages at GWAS SNP sites and calculate coarse-grained polygenic risk scores (PRSs; **Figure 1C**). In this way, we chart out a comprehensive exploration of how to quantify risks, and end with a mechanism to carefully sanitize datasets.

### Datasets and shared parameters

In some of the following sections, we test our tools using a series of simulations. The primary reference database employed is the 1000 Genomes Phase 3 database(Auton et al. 2015) (based on the human genome reference build *GRCh37*, fasta file *human_g1k_v37.fasta* (ftp.1000genomes.ebi.ac.uk/vol1/ftp/technical/reference), with 2,504 phased genotypes in total from 26 different sampled populations. The methods rely on the availability of phased reference genotypes, and currently work by reading through the chromosome-separated *vcf* files. We emphasize that, while we used the Phase 3 database based on the GRCh37 genome, the same method would carry over to GRCh38 or any other genome build. The only requirement is that the SNP positions in the query match up with those in the reference database, and the matching process proceeds in exactly the same manner. For all analyses, we filter out the low allele frequency SNPs, with minor allele frequency (MAF) restricted as: 0.05 ≤ *MAF* ≤ 0.5. The lower cutoff is a user-defined parameter in our program with a default value of 0.05. We choose such a cutoff with the intention of quantifying the leakage associated with even relatively common SNPs, as the identifying information contained in low MAF SNPs will be significant.

### Identification of individuals known to be within a database

We start with the simplest scenario, where an individual is known to be within a database. We tested out examples with different numbers of SNPs and varying values of the mutation rate/genotyping error (here we report the value of *λ* in the results, to simulate the effect of a particular genotyping error rate; see the *Methods* section for details on different mutation rate quantifications). For each of 5 mutation/error rates, we ran 10 iterations of the following simulation process. A single individual is selected at random from the full 1000 Genomes Phase 3 cohort of 2,504 individuals, and the genotype is simulated as described in the *Methods*. Given the set of 2,504 individuals, the chosen individual is likely to be different for each of the 10 iterations (even though sampling was done with replacement).

The variant selection procedure is repeated until we have *N*_*SNP*_ ∈ [1, …,40] for a particular simulation. To assess the HMM model, we consider for now only one chromosome (chromosome 17 was chosen at random for this analysis).

Therefore, for each error rate and iteration, we choose a single individual and simulate all 40 possibilities of *N*_*SNP*_ ∈ [1, …, 40] for that individual. The SNP selection is repeated from scratch for each of the 40 cases, so the SNP cohorts are not necessarily the same for the different values of *N*_*SNP*_. Also, note that it is possible to run the simulations with SNPs that are homozygous for the reference allele. This would yield the same trajectory prioritization in the HMM model. However, we attempt here to simulate cases where the focus is on identified alternate alleles, reported from either SNP beacons or noisy functional genomics data.

Finally, we find the mean and standard deviation across the 10 iterations of the minimum value of *N*_*SNP*_ for which a unique identification of the individual was made. However, in the presence of a mutation rate, it is possible for these unique identifications to be different from the ground truth samples. Therefore, we also provide the mean and standard deviation of the minimum value of *N*_*SNP*_ for which a correct and unique identification is made. For reference, we also provide the mean and standard deviation of the pooled list of MAFs across all 10 iterations for each mutation rate value. We present these results in **Table 1**.

**Table 1.** Results of the “individual known to be in the database” analysis, as a function of the scaled mutation rate *λ*, using *N*_*SNP*_ ∈ [1, …, 40] and 10 iterations (10 individuals chosen randomly with replacement) at each mutation rate. The columns shown are: the minimum number of SNPs required to obtain a unique identification of an individual, the minimum number of SNPs required for that identification to match the true identity of the sample, and the pooled minor allele frequency of all sampled SNPs across the 10 iterations at each mutation rate.

| Mutation/Error<br>Rate $\lambda = \frac{\theta}{2(N + \theta)}$ | Minimum number of<br>SNPs for unique ID,<br>$N_{SNP}^{Unique}$: Mean ± S.D.<br>(10 iterations) | Minimum number of<br>SNPs for correct ID,<br>$N_{SNP}^{Correct}$: Mean ±<br>S.D. (10 iterations) | Pooled MAF<br>distribution across all<br>10 iterations: Mean ±<br>S.D. |
| --- | --- | --- | --- |
| 0.0 | 6.7 ± 0.9 | 6.7 ± 0.9 | 0.25 ± 0.13 |
| 0.05 | 6.1 ± 2.0 | 9.9 ± 2.5 | 0.27 ± 0.13 |
| **0.1** | $7.5 \pm 2.0$ | $9.9 \pm 2.3$ | $0.25 \pm 0.13$ |
| **0.2** | $6.1 \pm 2.2$ | $21.6 \pm 7.9$ | $0.22 \pm 0.12$ |
| **0.3** | $6.2 \pm 1.9$ | $12.1 \pm 15.5$ | $0.22 \pm 0.13$ |

The results of the simulations indicate that, while an identification can be unique for nearly the same average 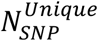 across all mutation rates, the ability to find the correct individual worsens with an increasing mutation rate. The mean value of 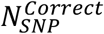 does not increase monotonically, but the standard deviation does. It is still noteworthy that unique identification can be made with a very small number of common SNPs (~6-8), with MAF values that are distributed fairly evenly across the range 0.05 ≤ *MAF* ≤ 0.5. Even in the presence of modest mutation rates (≲ 0.1), correct identification requires only about 10 common SNPs on average.

At the same time, this result may not be entirely surprising if one considers both the number of markers used in forensics studies, as well as the information content of a set of SNPs under Hardy-Weinberg equilibrium. With respect to the former, we note that the CODIS database expanded the number of core loci used from an original 13 to 20 in 2017(Federal Bureau of Investigation). Concerning the latter, it can be easily shown that with a set of independent SNPs under Hardy-Weinberg equilibrium and with MAFs in the same range as our analysis, a similar number of SNPs as shown here are sufficient to identify an individual in a database of the scale of 1000Genomes. Our purpose here is to confirm the power of a small set of SNPs, and to show that PLIGHT can evaluate the identifiability of an individual irrespective of the LD structure and MAFs of the query SNPs, and even in the presence of moderate amounts of noise.

### Identification of individuals from contaminated samples

To quantify the degree of leakage from samples that may have been in contact with multiple individuals, we chose to run a simulated experiment as follows. We (a) select five individuals; (b) draw 40 SNPs from a single chromosome for each of them (reference set of SNPs); (c) designate one individual as the primary “target” of the attack; (d) with a varying probability (ranging from 0 to 0.7), we randomly replace each SNP of the target individual with a SNP from one of the other four individuals (chosen with equal probability = 0.25); (e) using this “contaminated” SNP set, iteratively remove one SNP and run the inference procedure to determine if we can uniquely and correctly the target individual; (f) record the minimum number of SNPs allowing unique and correct identification. We run 3 iterations over the choice of individuals and associated reference SNP set, and 10 iterations where the contaminated set is randomly generated anew from each such reference set. The inference procedure used here (*PLIGHT_InRef*) operates under the assumption that the target individual is in the reference database (thus only the emission probabilities at each SNP matter, and not the transition probabilities). This experiment serves as a simulation of the case where a contacted object may have a primary user, but that contamination from other individuals may have occurred to an unknown degree. The replacement rate was also used as the error rate (*λ*) in the inference process (this is prior knowledge, but if unknown, the user would just iterate through several possible error rates in the inference procedure).

The results are shown in **Table 2**. We identify the number of cases of unique and correct identification of the target. For the pool of individuals, we considered two possibilities – the full 1000Genomes (1kG) set, and the CDX subpopulation – to explore the impact of the SNP diversity of the pool.

**Table 2.** Results of the contamination analysis, as a function of the replacement rate *λ*, using *N*_*SNP*_ ∈ [1, …, 40] and 30 iterations (3 iterations over reference SNPs, and 10 random contamination sets from the same set of reference SNPs). The columns shown are: the number of unique and correct identifications of the target individual (out of a total of 30 iterations), and the minimum number of SNPs for each successful run. These two columns are shown for both pools of individuals, the full 1kG set and the CDX set.

| Replacement Rate $\lambda$ | 1kG: Number of unique and correct IDs / total number of runs | 1kG: Average Minimum number of SNPs for correct ID across the successful iterations | 1kG: Average $\frac{\log(P_{HMM}) - \log(P_{GF})}{N_{SNPs}}$ | CDX: Number of unique and correct IDs / total number of runs | CDX: Average Minimum number of SNPs for correct ID across the successful iterations | CDX: Average $\frac{\log(P_{HMM}) - \log(P_{GF})}{N_{SNPs}}$ |
| --- | --- | --- | --- | --- | --- | --- |
| 0.0 | 30 / 30 | $7.0 \pm 1.3$ | $0.36 \pm 0.27$ | 30 / 30 | $7.3 \pm 1.4$ | $0.22 \pm 0.20$ |
| 0.1 | 30 / 30 | $8.2 \pm 2.4$ | $0.33 \pm 0.20$ | 30 / 30 | $7.5 \pm 2.0$ | $0.29 \pm 0.13$ |
| 0.2 | 30 / 30 | $13.9 \pm 5.8$ | $0.44 \pm 0.14$ | 30 / 30 | $10.8 \pm 5.6$ | $0.32 \pm 0.15$ |
| 0.3 | 20 / 30 | $16.8 \pm 8.8$ | $0.46 \pm 0.20$ | 26 / 30 | $14.8 \pm 7.4$ | $0.39 \pm 0.13$ |
| 0.4 | 4 / 30 | $13.8 \pm 6.5$ | $0.63 \pm 0.23$ | 7 / 30 | $11.6 \pm 3.9$ | $0.41 \pm 0.09$ |
| 0.5 | 0 / 30 | N/A | N/A | 0 / 30 | N/A | N/A |
| 0.6 | 6 / 30 | $26.8 \pm 7.0$ | $-0.20 \pm 0.09$ | 0 / 30 | N/A | N/A |
| 0.7 | 22 / 30 | $23.5 \pm 8.2$ | $-0.35 \pm 0.23$ | 1 / 30 | $36.0 \pm$<br>N/A | $-0.07 \pm$<br>N/A |

We first note the expected behavior that, with increasing replacement rates, the average minimum number of SNPs required for correct identification of the target individual increase. However, we also find an interesting trend for the 1kG pool, where the number of successful identifications drops to 0, before increasing again at higher error rates. We do not see a similar trend with the CDX population. Given that we explicitly chose the target individual’s SNPs to be heterozygous or homozygous in the low MAF alternate allele, we surmise the following. Against a background of the full 1kG set, replacing more SNPs in the target SNP set results in more homozygous reference alleles being introduced. This may increase the similarity to other individuals in the 1000Genomes reference database, and thereby result in more failures to detect the target individual. However, upon further increasing the number of replacements from four other individuals, as well as the error rate in the inference procedure (see above), we suspect that the similarity to any other individual in the database drops again, leaving the target individual as the most likely result. In the case of the CDX individuals, trading SNPs between relatively similar individuals effectively masks the target individual above a certain replacement rate. Thus, the result supports the logical notion that it is easier to protect a single individual from reidentification in the midst of genetically similar individuals.

We also utilized PLIGHT’s output of different genotype probabilities to demonstrate the degree to which the SNPs were independent. In this example, we used the per-SNP difference between the logarithm of the joint probability of the SNPs under the HMM model (*log*(*P*_*HMM*_)), and the probability under the assumption of independent SNPs. For the latter, we considered the genotype frequencies (*log*(*P*_*GF*_)) rather than the MAFs under Hardy-Weinberg equilibrium (though both are output by PLIGHT) as *PLIGHT_InRef* focuses on matching genotypes and not haplotypes. We see that the *log*(*P*_*HMM*_) > *log*(*P*_*GF*_) on average for low mutation rates, but this trend reverses at higher mutation rates. At low mutation rates, the non-independence of SNPs would lead to a higher *log*(*P*_*HMM*_), while at high mutation rates the emission probabilities deviate from 1 sufficiently to lower *log*(*P*_*HMM*_) relative to *log*(*P*_*GF*_) (see definitions in *Methods* section). Interestingly, for a small number of runs, a small negative 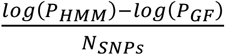 was observed even with 0 replacement rate (**Supplemental File S1**). This is likely due to round-off errors.

### Mosaic overlap between query individuals and database individuals

We next evaluated the performance of the program on *mosaic* individuals, i.e. individuals whose genome is constructed by sampling the diploid genomes of two or more source individuals, using simulated analyses (the mutational process is the same as in the previous section). A pair of individuals is chosen at random from the 1000 Genomes set of individuals. The diploid mosaic genome is constructed for *N*_*Chr*_ sampled chromosomes using the simulation scheme described in the *Methods* section.

#### Exact search within a reference database of 400 haplotypes

For the first example, we use the *PLIGHT_Exact* module employing the full Li-Stephens model. Two individuals were selected, and the first half of the SNP genotypes were taken from one individual and the other half from the other. The mutation rate was set to 0, while the fixed per base recombination rate *c*_*l*_ was set at 0.5 cM/Mb and the default linear recombination model was used. The choice of 0.5 cM/Mb was chosen close to the value of 0.4 cM/Mb, a biologically plausible average rate of recombination used in previous HMM studies (Li and Stephens 2003; Howie et al. 2009). This value led to reasonable exploration of the haplotype space, without devolving into a uniform consideration of all haplotypes as equally probable. To understand this, consider the two extremes: a very low value of *c*_*l*_ will prefer to elongate the same haplotypes for the entire length of the observed SNP positions without crossing over, i.e. the emission probabilities will dominate in the overall likelihood; a very high value will lower the barrier to cross-overs, causing the probabilities to be equal across all choices of haplotypes, i.e. the transition probabilities will dominate in the overall likelihood.

To limit the dimensionality of the matrices used in the exact case, we only search for matches among 200 individuals (= 400 haplotypes) from the reference database. We sampled *N*_*SNP*_ = 30 SNPs from each of *N*_*Chr*_ = 3 chromosomes (chromosomes 1, 2 and 21). The two sampled individuals were HG00360 (first half) and HG00342 (second half). Note that, given the choice of random parameters in the SNP selection process for the simulated query genomes, the total length of the query genomes for each of the chromosomes was much smaller than the length of the chromosomes: in the following example, the SNP positions ranged from 1.03 Mb to 37.3 Mb for chromosome 1, from 0.4 Mb to 22.1 Mb for chromosome 2, and from 14.7 Mb to 38.4 Mb for chromosome 21. The results are provided in **Figure 3** and **Supplemental Fig S2**.

**Figure 3.**
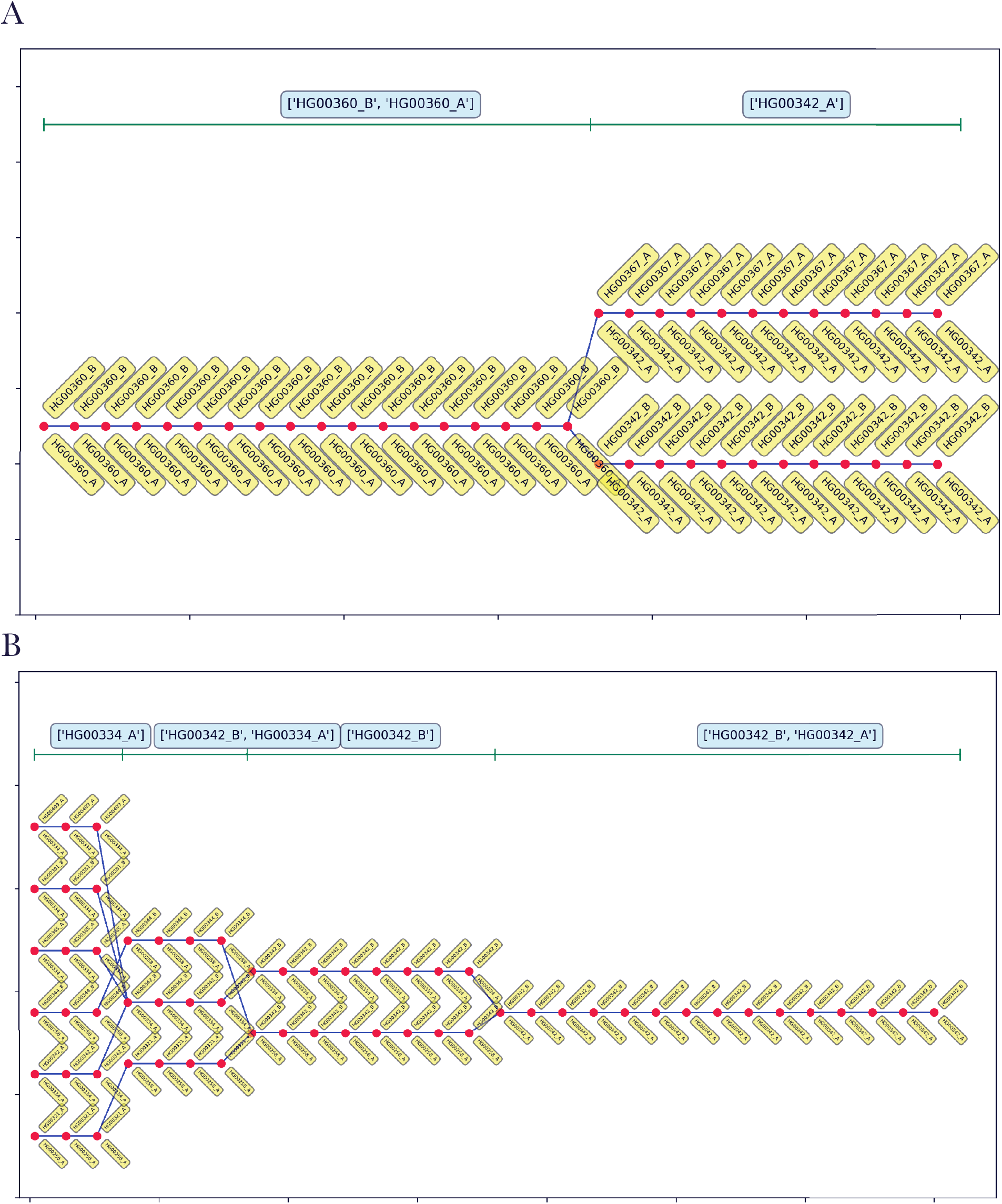
Best-fit genotypic trajectories from PLIGHT_Exact for the diploid mosaic genome of HG00360+HG00342 constructed across 30 SNPs each for chromosomes 1 and 2 (results for chromosome 21 shown in **Supplemental Fig S2**). The composition of the best-fit pair of haplotypes at each locus is depicted by two yellow tags, one below and one above the red dots. (A) Trajectory for chromosome 1. (B) Trajectory for chromosome 2.

**Figure 3A** shows the genotypic trajectories for chromosome 1, and **Figure 3B** shows the same for chromosome 2 (the results for chromosome 21 are provided as **Supplemental Fig S2**). The labeling scheme we use involves the splitting of the phased genotypes in the reference database into the two component parental haplotypes, with the (arbitrary) haplotype labels “A” and “B” appended to the name of the reference individual. The labels for the pair of haplotypes in the best-fit trajectories are depicted by one yellow tag below and another above the red dot marking each locus of each trajectory. The results for chromosome 1 indicate that the correct mosaic was identified as one of the two genotypic trajectories. The second trajectory consists of a mixture of one haplotype from the true individual (HG00342) and the other from a different individual (HG00367). The two trajectories branch out from HG00360 at the same SNP, indicating that for the last set of SNPs, HG00367_A and HG00342_B are likely identical (there are no noise mutations introduced in this particular simulation). Indeed, the vcf files confirm this to be the case, at least for the last 13 SNPs in the simulation. The optimal haplotypes for certain stretches of the chromosome are identified at the top of the figures, with the boundaries between the stretches marked by green ticks. These optima are calculated by simply maximizing the frequency of occurrence of the haplotypes in these regions. An important point to note here is that even though the simulation drew the last 14 SNPs from HG00342, the transition from HG00360 to HG00342 in the solution occurs only for the last 12 SNPs. This is because the trajectories require a certain number of additional genotypic steps to build enough probability to warrant a transition. If the inference process is seen as a combined approach to identify the stretches of best-fit genotypes, as well as the best-fit boundaries between these stretches, then this method does lead to a certain fuzziness in the identification of the boundaries. However, an uncertainty is likely to exist in any dataset, regardless of the method, if there is even a little error in the genotyping or any mutations.

The trajectories for chromosome 2 (**Fig. 3B**), on the other hand, include HG00342, but not HG00360. Several other trajectories and branch points occur in the region of the true HG00360 segment. The reason for the ground truth sample not arising as a part of the solution is likely that, at the current rate of recombination, it is more likely for one of the haplotypes (HG00342_B) to be maintained across a longer stretch of the chromosome, which then results in alternative haplotypes being selected over shorter segments than the true HG00360 segment. To test whether this may be the case, we ran the same sample through a calculation with double the previous recombination rate (= 1 cM/Mb). The results (**Figs. S3A, B and C**) indicate that increasing the recombination rate does indeed cause the true HG00360 segment to be included in the solution. The increase in the recombination rate is more favorable to the transition from one haplotype to another and thus allows for a greater exploration of the haplotype space. This is also evident from the increase in the average number of best-fit haplotypes per SNP evident from **Fig. S3**. The results for chromosome 21 (**Fig. S2**) do include the true individual HG00360, in addition to several alternative trajectories.

#### Truncated algorithm

We subsequently ran the same SNP set through the truncated algorithm to assess the degree of compressibility of the trajectories and the potential for memory reduction. The results, which indicate a considerable degree of compressibility of the best-fit trajectories, are presented in the Supplemental Results and **Supplemental Fig S4**.

#### Iterative and approximate search within a reference database of 5,008 haplotypes

The *PLIGHT_Iterative* algorithm contends with the memory requirements of a large reference database by subdividing the database into subgroups and running the HMM on the smaller haplotype cohorts. Each round of subdivision is run several times, with different haplotypes assigned to different subgroups each time. We test the same synthetic input dataset as for *PLIGHT_Exact*, but now search through the entire 1000 Genomes phase 3 database with 5,008 haplotypes. We ran two replicates for each of two sets of parameters: number of iterations (a) *n*_*iter*_ = 20; and (b) *n*_*iter*_ = 30; the subgroup size, *S*_*sg*_, is chosen to be *S*_*sg*_ = 300, and the recombination rate set to *c*_*l*_ = 0.5 cM/Mb. The randomness of the algorithm is apparent from the fact that there is no unequivocal improvement in the identification of component individuals in going from *n*_*iter*_ = 20 to *n*_*iter*_ = 30. While it is possible that a significantly larger increase will consistently improve the mixing of the haplotypes in general, these runs are mostly able to identify the true component individuals in the mosaic sample, especially when information across the chromosomes was combined. Accordingly, the default value in the algorithm is set to *n*_*iter*_ = 20. The *consensus* trajectories, which are trajectories containing haplotypes frequently observed across chromosomes (see *Methods*), include both HG00342 and HG00360 in three out of four of the runs (though not necessarily within the same trajectory). Additionally, the chosen SNPs on each chromosome have different capacities for discerning the ground truth. The consensus results for chromosomes 1 (one set of replicates in **Figure 4** and the other in **Supplemental Fig S5**) and 21 (**Supplemental Fig S6**) include the true HG00360+HG00342 combination within the same trajectory for one and two out of the four runs, respectively.

**Figure 4.**
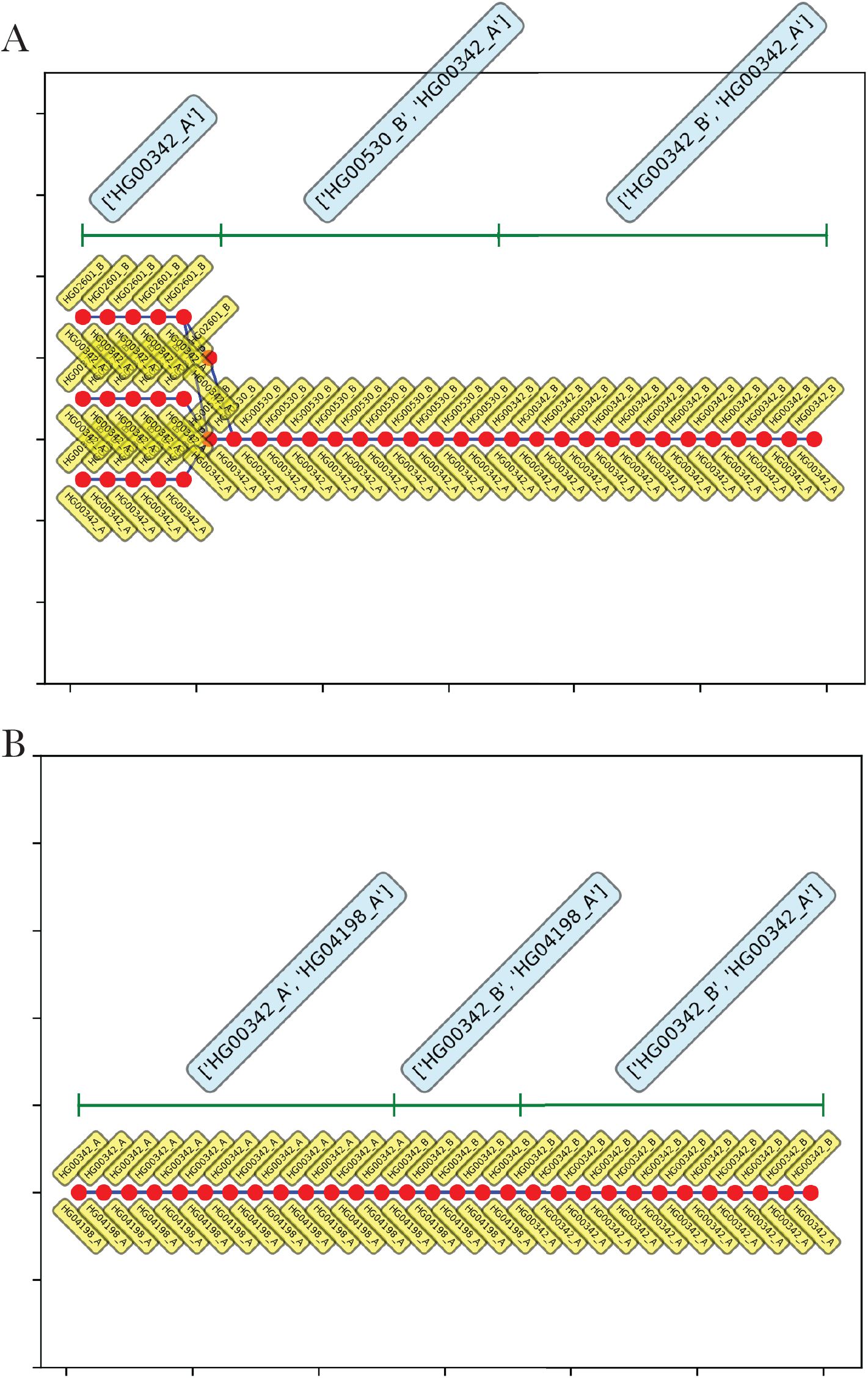
Consensus genotypic trajectories from PLIGHT_Iterative for the diploid mosaic genome of HG00360+HG00342 constructed across 30 SNPs in chromosome 1, where the consensus score is evaluated by weighting the haplotypes in each trajectory in proportion to their occurrence across all three chromosomes. The composition of the best-fit pair of haplotypes at each locus is depicted by two yellow tags, one below and one above the red dots. (A) n_iter_ = 20, replicate 1; (B) n_iter_ = 30, replicate 1. Shown at the top of each panel are the most frequent haplotypes within each segment indicated. The second replicates of n_iter_ = 20 and n_iter_ = 30 are shown in the Supplemental Materials. Thus, the iterative, mixing algorithm is able to pick out the ground-truth component individuals in the best-fit trajectories, albeit with a certain degree of randomness.

Overall, the results in this section demonstrate that it is possible to identify, with high confidence, the individuals contributing to local segments of the genome. This is important to demonstrate as one of the main strengths of PLIGHT lies in not just identifying a one-to-one match between a query set and a specific individual in the database, but potential overlaps in local regions of the genome which may contribute to privacy leakages. In the following sections, we explore ways in which this local information may lead to privacy breaches.

### Kinship analysis

Having analyzed synthetic mosaics in the previous results, we seek to benchmark the methods on known, *natural* mosaics in the form of a kinship analysis. We use a set of 13 individuals taken from the related samples cohort of the 1000 Genomes Phase 3 (ftp://ftp.1000genomes.ebi.ac.uk/vol1/ftp/release/20130502/supporting/related_samples_vcf/related_samples_panel.20140910.ALL.panel) to link them to those individuals among the Phase 3 main release cohort of 2,504 phased genotypes that are stated to be first-(parents, children or siblings), second- or third-order relatives (as provided in an associated pedigree file (ftp://ftp.1000genomes.ebi.ac.uk/vol1/ftp/technical/working/20130606_sample_info/20130606_g1k.ped). For this study, we choose a single chromosome for each individual with the parameters: (I) *N*_*SNP*_ = 20, *n*_*iter*_ = 20; and (II) *N*_*SNP*_ = 30, *n*_*iter*_ = 30; the subgroup size was chosen to be *S*_*sg*_ = 300, and the recombination rate set to *c*_*l*_ = 0.5 cM/Mb. The chromosomes were chosen at random, and are different in general between cases (I) and (II). A successful identification is indicated as the inclusion of the related individual anywhere within the best-fit trajectories. The results are presented in **Table 3**.

**Table 3.**
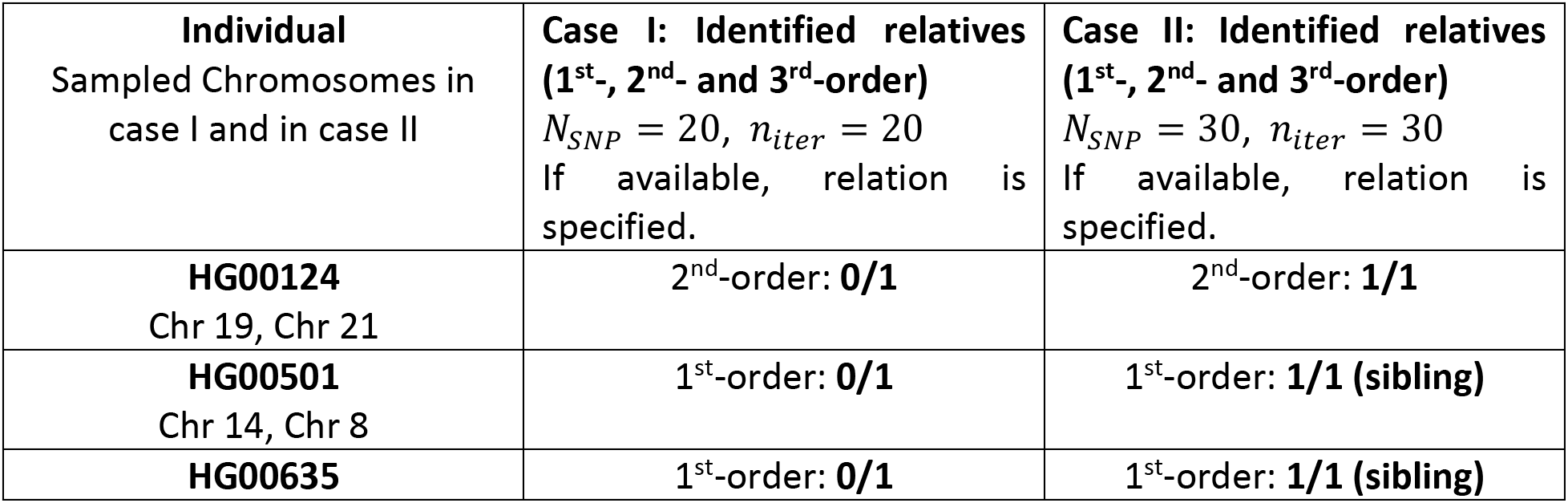
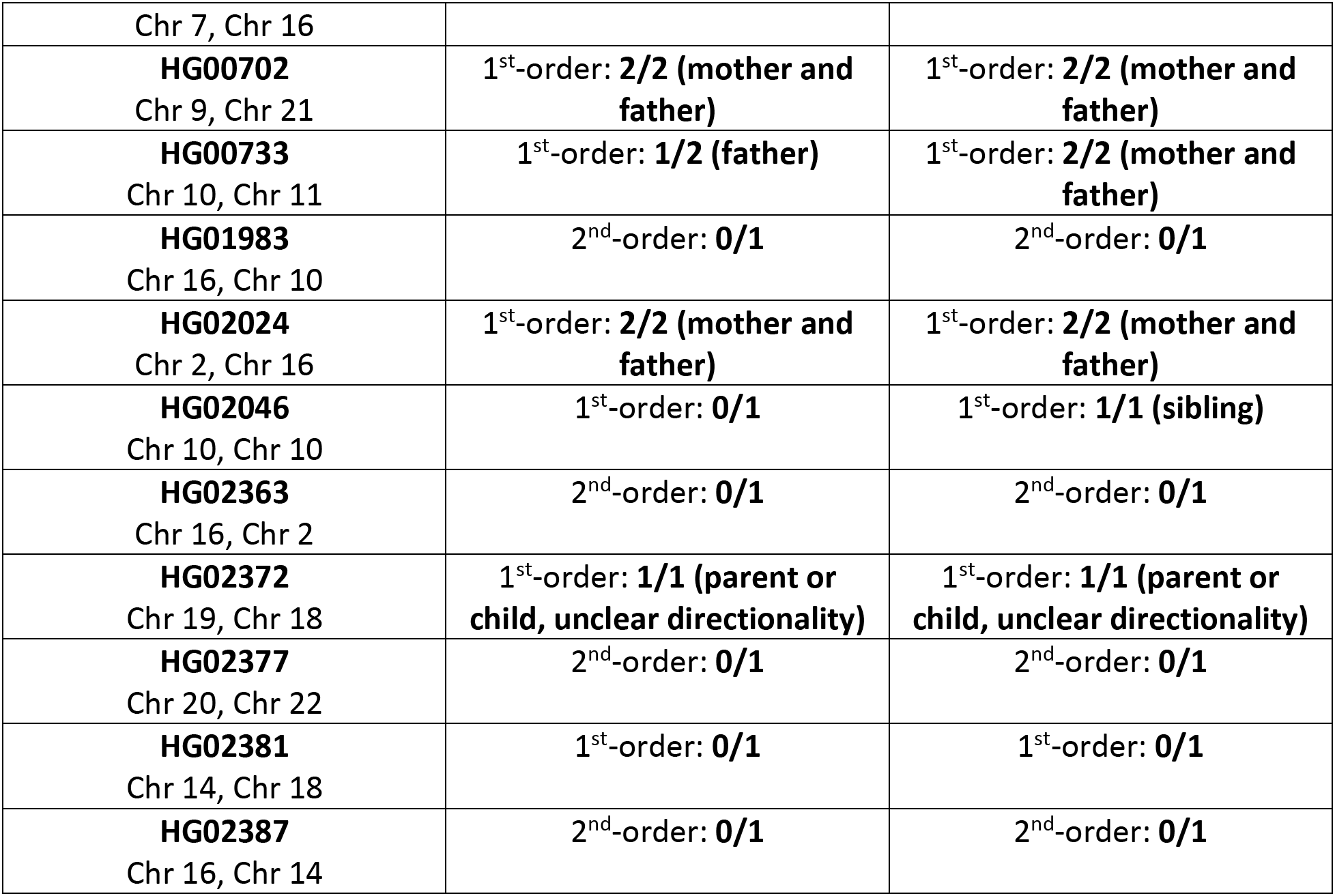
Results of the kinship analysis for the two sets of parameters. Case I: *N*_*SNP*_ = 20, *n*_*iter*_ = 20; and Case 2: *N*_*SNP*_ = 30, *n*_*iter*_ = 30. The degree of relatedness is indicated as “1^st^-, 2^nd^- and 3^rd^-order” and when a successful identification is made, the relationships of the identified individuals to the query individuals are explicitly specified, if available from the pedigree file (ftp://ftp.1000genomes.ebi.ac.uk/vol1/ftp/technical/working/20130606_sample_info/20130606_g1k.ped).

The analyses were designed to simultaneously explore the impact of several parametric factors in the identification of relatives: (a) *n*_*iter*_, (b) *N*_*SNP*_ and (c) chromosome identity. From the results in preceding section, it is likely that changing *n*_*iter*_ had minimal impact. At the same time, previous results also indicated that the choice of SNPs and which chromosome they occur on will impact the best-fit trajectories. To parse these two effects, we note that irrespective of the query individual and the chromosomal choice, increasing the SNPs clearly improves the efficiency of the kinship discovery (comparing the 2^nd^ and 3^rd^ columns of Table 3). While for several queries no identification was made for either set of parameters, there is a clear and understandable increase in risk for 1^st^-order relatives compared to 2^nd^-order relatives. There was only a single instance in which a 2^nd^-order relative was discovered for a query individual, and in that instance there were no 1^st^-order kin in the 1000 Genomes main cohort to confound the identification process.

However, even in the case of 1^st^-order relatives, the identification process is not perfect. For example, the individual HG02381 has a sibling in the 1000Genomes main release cohort. This sibling was not discovered even after increasing the number of SNPs to 30. The randomness of *PLIGHT_Iterative* might contribute to this, beyond the inherent informativeness of the SNP set. To test this out, we separated out the 93 individuals belonging to the same population group and ran the query SNPs through *PLIGHT_Exact*. In this more restricted case, one of the haplotypes of the sibling was found as a match, but only for a part of the trajectory. This example is therefore a case where the genetic relative remained concealed due to a combination of the inherent SNP informativeness (as evidenced by a lack of kin identification even on a smaller, exact run), size of the database considered, and the approximation of the algorithm.

In general, we show that even a modest number of common SNPs have the capacity to reveal genetic relatives, especially 1^st^-order relatives, of query individuals in a cohort of size ~5,000 haplotypes. We did not carry out a comprehensive search for the minimum number of SNPs to identify relatives because (a) *PLIGHT_Iterative* is more computationally intensive and iterating over multiple query sets for many individuals can become unfeasible, and (b) without prior knowledge on which genomic regions overlap between relatives, a change in the number of randomly selected SNPs may not yield clear improvements. Our primary goal, however, is to show that even small SNP sets can reveal close relatedness.

### Inference based on SNPs obtained from coffee cups

Moving away from simulated examples based on the reference database itself, we use a set of SNPs that were obtained from DNA samples acquired from swabs of used coffee cups in ref. (Gürsoy et al. 2020) (more details in *Methods*). As described in ref. (Gürsoy et al. 2020), surreptitiously acquired samples such as these pose significant identification risks to individuals. For our purposes, they additionally represent a source of noisy (i.e. error-prone) SNPs that can be used to test the methods herein.

We filter the SNPs obtained from the coffee cups by genotyping quality and read depth (‘GQ > 99 & DP > 10’ chosen as the vcf parameter filters, applied using *bcftools* version 1.10.2), to increase the confidence in the choice of genotypes. We subsequently select 30 SNPs each from chromosomes 3 (~position 4 Mb to 173 Mb) and 6 (~position 11 Mb to 166 Mb). We compare these against the blood-tissue-derived genotypes as the gold-standard, as well as to the 1000 Genomes phase 3 vcf files. All 30 chromosome 3 SNPs and 29 chromosome 6 SNPs (1 missing) were found in the blood-tissue and 1000 Genomes vcf files.

We checked the 60 SNPs obtained from the coffee cups against the true genotypes of the query individual: 8 genotypes out of 30 for chromosome 3 were incorrect, while 4 genotypes out of 29 were incorrect for chromosome 6; all incorrect cases involved calling homozygous alternate as opposed to the correct heterozygous genotypes. The incorrect calls are likely to arise due to possible contamination of the coffee cups by DNA from other individuals, as well as due to the inherent noisiness of genotypes obtained from the coffee cups. Accordingly, we include non-zero mutation rates in our inference to account for errors or contamination. Also, we note that both the coffee cup genotypes and the blood tissue gold-standard genotypes are unphased: we distribute the SNP dosages in the blood tissue sample based on positional occurrence in the called genotype; thus, for a genotype call of “0/1”, “0” is assigned to the first haplotype and “1” to the second.

We carry out two analyses. The first involves merging the gold-standard SNP set with the 1000 Genomes Phase 3 vcf file, while making sure that only overlapping SNPs were retained. We subsequently determine whether ~30 SNPs each on chromosomes 3 and 6 would be sufficient to identify the true query individual, using both *PLIGHT_InRef* (where a simpler algorithm is run with the recombination rate set to 0) and several runs of *PLIGHT_Iterative*.

#### With 30 SNPs: Query individual in the reference database

The *PLIGHT_InRef* algorithm, with a mutation rate *λ* = 0.2, correctly and uniquely identifies the query individual out of the full set of 2,504 genotypes for both query chromosomes separately. We ran then four inference runs using *PLIGHT_Iterative*, where the full set of 5,008 haplotypes was used as the reference database: (a) *S*_*sg*_ = 200, mutation rate *λ* = 0.1; (b) *S*_*sg*_ = 200, mutation rate *λ* = 0.2; (c) *S*_*sg*_ = 300, mutation rate *λ* = 0.1; (d) *S*_*sg*_ = 300, mutation rate *λ* = 0.2. The variation of the subgroup size parameter was done to determine how stable the results were to different bootstrap sample sizes. In every single case, we detect one of the two haplotypes of the correct query individual, while the other inferred haplotype is drawn from the 1000 Genomes set. The lack of both query haplotypes showing up in this instance is likely due to a combination of the high error rate in the coffee cup SNP set and the approximation inherent in the iterative algorithm. We explored the reasons for the consistent appearance of the same gold-standard query haplotype in all the searches, and found that since all the errors involved incorrectly calling a genotype of “1” as a “2”, and since all unphased heterozygous genotypes in the blood tissue sample were called as “0/1”, the errors in the coffee cup SNPs would be associated with the first haplotype in our arbitrary phasing scheme (see above). Thus, the algorithm finds the second gold-standard query haplotype consistently, as it correctly aligns with the coffee cup SNPs.

#### With 30 SNPs: Query individual not in the reference database, and coarse-grained imputation

The second analysis on the coffee-cup-based SNPs involves running *PLIGHT_Iterative* on the query SNPs to search through the 1000 Genomes Phase 3 reference database without including the true query individual’s genome. We ran the same four combinations of parameters as in the preceding analysis. For both chromosomes, there are haplotypes and haplotype combinations that are found robustly in multiple runs, indicating the algorithm is able to find close matches to the query SNPs. In subsequent analyses, we used some of these best-fit trajectories to explore other instances of inference.

We take trajectories from one of the runs (the results with *S*_*sg*_ = 200, mutation rate *λ* = 0.2) and reconstruct the mosaic genomes (see *Methods*). We use the overlapping SNPs between the gold-standard query vcf and the 1000 Genomes phase 3 reference vcf to determine the total fraction of matching genotypes, and the correspondence score *C*, a weighted fraction that accounts for the rarity of each SNP in the reference database, which would influence the likelihood of observing a match by random chance (see *Methods*).

We also calculate background values for 99 randomly chosen individual genomes from the 1000 Genomes dataset, ensuring that there was no overlap with the mosaic genome haplotypes. There are three matching trajectories for chromosome 3 and one trajectory for chromosome 6. The exact-matching-genotype fractions for the trajectories in chromosome 3 are 0.43, 0.43 and 0.43, compared to the background matching fractions ranging between 0.36-0.44 [Welch’s two-sample t-test (for a null hypothesis of both distributions being the same) p-value = 2e-10]. For chromosome 6, the matching fraction is 0.40, with a background between 0.34-0.44. The correspondence scores are: chromosome 3, *C* = 0.32, 0.32, 0.32 [background 0.28-0.33, p-value = 0.0002]; chromosome 6, *C* = 0.31 [background 0.29-0.33]. Overall, while there is some evidence for correct imputation of the intervening genomic regions between observed SNPs, we sought to confirm the results, since the few samples drawn still fall within the background ranges.

#### With 90 SNPs: Query individual not in the reference database, and coarse-grained imputation

Given the results above, we explore whether the imputation process improves with a larger number of SNPs. We select an additional 60 coffee cup SNPs from chromosomes 3 and 6 for further analysis: for chromosome 3, all 90 are found in the gold-standard blood-tissue genotypes, 89 in 1000 Genomes Phase 3; for chromosome 6, 88 are in the gold-standard blood-tissue genotypes and in 1000 Genomes. We conduct all the above tests for this extended set of ~90 query SNPs. Running *PLIGHT_Iterative* on the in-reference case allows correct identification of both haplotypes of the query individual for chromosome 6, while only one correct haplotype is found for chromosome 3 (we did not run the exact inference, as 30 SNPs was clearly sufficient to find the correct query individual). While some haplotypes occur in both the 30-SNP and 90-SNP runs, the resulting trajectories are different in general (i.e. different number and haplotype composition). We subsequently use one of the trajectories from the runs where the true query individual’s genome is not in the reference database (the results with *S*_*sg*_ = 300, mutation rate *λ* = 0.1). There are 8 best-fit trajectories for chromosome 3 and 1 for chromosome 6. Comparisons to a background of 98 individuals are as follows: The exact-matching-genotype fractions for the trajectories in chromosome 3 are 0.45, 0.44, 0.44, 0.44, 0.43, 0.43, 0.43 and 0.44, compared to the background matching fractions ranging between 0.36-0.46 [Welch’s two-sample t-test p-value = 2e-8]. For chromosome 6, the matching fraction is 0.43, with a background between 0.34-0.43. The correspondence scores are: chromosome 3, *C* = 0.33, 0.33, 0.33, 0.33, 0.31, 0.31, 0.32 and 0.33 [background 0.28-0.33, p-value = 0.0001]; chromosome 6, *C* = 0.33 [background 0.29-0.33]. Overall, while the degree of matching improves a little in going from the 30-SNP case to the 90-SNP case, the change is small. We note that the comparisons here reflect the results of matching of genotypes across entire chromosomes. However, we imagine that within any LD blocks associated with the query SNPs we would observe better matching on average than the global results demonstrated here. We reiterated the analysis with a 300-SNP query set as well, and found similar results: the significance of the genotype matching increases for the mosaic trajectories relative to the background.

In summary, it was extremely easy to identify an individual from amongst 5,008 haplotypes, using just 30 SNPs on a single chromosome, even in the presence of significant errors. Searching against individuals in a reference database that does not include the query individual also yields best-fit trajectories that are robustly found by *PLIGHT_Iterative* across different parameter settings. Expanding to a larger set of query SNPs indicates that the haplotype composition of the trajectories does change to reflect the new query SNPs, but with some haplotypes stably appearing in both sets of runs. The coarse-grained imputation at the level of 30 and 90 SNPs hints at only subtle differences relative to the background. We imagine that these differences would be amplified in the case of a much larger set of SNPs, and when focusing on local regions of the genome in the vicinity of the query SNPs.

### Predicting genotypes at GWAS loci and polygenic risk score (PRS) analysis

The coarse-grained imputation we carried out in the previous section hints at the possibility that the trajectories inferred may leak more information about intermediate regions of the genome. This would undoubtedly be the case for SNPs in LD with the query SNPs, but if longer distance correlations in SNP occurrences exist, there would be added risk from such correlated SNPs. If this were the case, even partial matches (i.e. matches in specific regions of the genome, without overall identification of the query individual) to the query SNPs may allow an attacker to infer local information, say, the risk of certain genetic diseases.

To evaluate this risk, we probe scenarios where either excessive mutations or the absence of the individual in the reference set may prevent the identification of the true individual, but the set of best-fit diploid mosaic genomes may collectively provide hints about the true individual. We approximately calculate linear polygenic risk scores (PRSs) of these individuals for all the phenotypes in the GWAS catalog version 1.0.2 (Buniello et al. 2019) to see if we could infer certain phenotypes that are outliers for the set of alternate individuals. We analyze one of the runs in the previous section, where the query individual was a mosaic of HG00360 (first half) and HG00342 (second half), and the best-fit trajectories were identified using *PLIGHT_Iterative* with *n*_*iter*_ = 20 and *S*_*sg*_ = 300. There were 8 parallel trajectories for chromosome 1, 15 for chromosome 2, and 2 for chromosome 21. The PRS calculation was carried out on all these trajectories independently and the PRSs across all trajectories (for each chromosome separately) for each trait were compared with the PRSs for a 50% HG00360 / 50% HG00342 mosaic as the true sample. Four different statistical measures were evaluated as described in the *Methods* section. We note that the scenario described for the simulated example is slightly artificial, as some of the best-fit trajectories included the ground-truth individuals as well. Removal of these trajectories was not an option in this case, given the pervasiveness of the true individuals in the best-fit results. However, we discuss the calculations here as an illustrative example of the sort of aggregative attack that may be carried out. The results of the calculation are presented in **Table S2**.

We performed the same analysis for the coffee cup SNPs as well, evaluating the 30- and 90-SNP cases (**Table S3**), as well as the 300-SNP case (**Table S4**). For the simulated and coffee cup PRS score comparisons, we mostly see a high cosine similarity between the true and mosaic reconstruction PRSs. However, the corresponding cosine similarities for the background individuals are similarly high. There is also noise in these metrics for all comparison cases, resulting in some negative or low cosine similarities. Therefore, while there are hints that some degree of aggregative leakage of information may be happening, the evidence in these particular examples is not very strong.

Instead, we sought to demonstrate the potential risks by constructing an artificial example. We construct a query set of 90 SNPs where the selected SNPs were within ±2 kb of known GWAS SNPs for the phenotype “Height”, and which should have a higher chance of being in LD with GWAS variants. We choose “Height” as a phenotype due to the abundance of GWAS variants, as well as it being a more innocuous phenotype to consider in this demonstration. In the windows considered for these SNPs, 26 GWAS variants are found for each chromosome. We ran *PLIGHT_Iterative* with *n*_*iter*_ = 20 and *S*_*sg*_ = 300 on the query set, and study the resulting set of 7 trajectories for chromosome 3 and 1 trajectory for chromosome 6. First, we did a direct comparison of how well the inferred trajectories impute unseen GWAS SNPs. That is, we looked at the 22 SNPs in each chromosome that are not directly in the query set and checked how well the inferred trajectories have matched the query genotypes. For chromosome 3, the 7 trajectories produced {14, 14, 14, 14, 14,11, 11} matches out of 22 SNPs. We also checked the same SNP sites for 100 randomly sampled individuals, and ran a Welch’s two-sided, two-sample t-test between the two distributions, obtaining a p-value of 2.2e-5. However, a couple of the background samples matched the GWAS SNPs to a higher degree (15 and 16 SNPs out of 22). We therefore looked at the consensus across the trajectories and found that the 7 trajectories matched the query genotypes perfectly at 10 GWAS SNP sites, were off by a dosage of 1 at 7 sites, and had mixed results for the remaining 5. The background individuals had mixed results across all SNP sites. The one trajectory for chromosome 6 yielded a match rate of 16 out of 22 and a two-sided, one-sample t-test with respect to the background distribution for the same 100 individuals yielded a p-value of 2.2e-16. None of the background individuals did better than 16 out of 22 matches. Thus, while the imputation of GWAS SNPs was not perfect, there are certain loci where the inferred trajectories do better at imputing the GWAS SNPs. We can foresee cases where an attacker might apply such an approach to phenotypes such as eye color, which are associated with a relatively small number of SNPs(Meyer et al. 2020).

We also assessed whether the inferred PRSs are a better match than a background set of PRSs from 100 randomly sampled individuals. We consider the statistical significance of the deviation from the true sample’s PRS (a Welch’s two-sided, two-sample t-test for chromosome 3, and a two-sided one-sample t-test for chromosome 6). The PRS deviations are resulting p-values are: chromosome 3 = 2.5e-6, and chromosome 6 = 0.008. In both cases, the trajectory-based PRSs were closer to the true sample’s PRS than the mean of the background PRSs (distributions show in **Supplemental Fig S7**). Here, the number of trajectories is small, and the example was purposely constructed to bring out LD effects, but we believe that the results are indicative of some information leakage.

Our intention in presenting this result is to demonstrate the *types* of privacy incursions that may be conducted in the future by utilizing partial inference. While the cross-phenotype analysis was inconclusive, the targeted analysis indicates that there exists the potential for inferred trajectories to leak information about certain SNP dosages and phenotypes. This information leakage is likely to be stronger when query SNPs are in the vicinity of GWAS variants for the phenotype of concern. An attacker may query genomic regions proximal to disease loci, and even specifically amplify proximal genetic material.

### Sanitization of SNPs based on Inferred Trajectories

The preceding sections demonstrated the various ways in which PLIGHT can be employed to determine the types of privacy risks associated with a SNP set. We also designed an algorithm to use the inferred trajectories to inform the removal of individual SNPs from a dataset, called *PLIGHT_SanitizeGenotypes*. Sanitization would be applied for those datasets whose primary purpose is different from the direct reporting of variants, but from which variants can be inferred. Functional genomics assays are prime examples of such datasets, as well as environmental samples where the identity of any of the contributing individuals needs to be protected. The algorithm is a modification of the simple strategy of removing SNPs based on rarity, i.e. based on the MAF. The subset of SNPs with the fewest parallel best-fit trajectories are identified. These are “bottleneck” positions in the query set where the number of parallel trajectories passing through them is the smallest, and so represent the maximum information about the underlying identity of the query individual. Specifically, the number of trajectories passing through a locus is quantified by the number of unique haplotype pairs (*N*_*UHP*_) at that locus (**Supplemental Fig S8** and defined in the Methods section). Often, multiple SNPs have the same number of trajectories passing through them, and so the SNP with the lowest MAF in this subset is removed. We emphasize that this is just one strategy among many possibilities, and depending on the use case, a user may choose to make other sanitization decisions based on the output of PLIGHT.

To demonstrate the use of *PLIGHT_SanitizeGenotypes*, we construct 10 mosaic SNP sets consisting of two source individuals, with half of the 30 SNPs drawn from one and the remaining from the other. We proceed to remove one SNP at a time, either based on the PLIGHT strategy or on the simpler MAF approach, and infer the best-fit trajectories on the new, sanitized SNP set. The results are compared to identify the relative merits of the two sanitization strategies.

The simplest way of assessing whether a sanitization strategy works is by calculating the degree to which the true source individual is hidden among several others (the source individual will never completely disappear in a noise-free situation). Also, we have the opportunity of searching for the source individual globally over the whole query set, or locally at particular sites. Accordingly, we settled on three metrics (formal definitions in the *Methods* section): (1) the total informational entropy of the identified individuals, *S*_*Ind*_; (2) the maximum probability of observing any of the two source individuals in the trajectory, 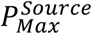; and (3) the per-SNP entropy of identified individuals, if either of the source individuals is found at that SNP, *S*_*Per—SNP*_ (Supplemental Fig S8). All three metrics are calculated as a function of the number of SNPs removed. We note that for each of the ten examples, the number of SNPs removed is different as we stopped the removal process for each example once the number of inferred trajectories grew very large.

We find from the plots of *S*_*Ind*_ (**Supplemental Fig S9**) and 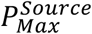 (**Supplemental Fig S10**) that there is no pattern indicating that PLIGHT does a better job at increasing overall entropy, or reduces the probability of detecting the source individuals across all SNPs. Both these sets of plots evince considerable variability across runs. On the contrary, the plots of *S*_*Per—SNP*_(*i*) (**Figures S11** and **S12**) show a more consistent increase in the entropy per SNP for PLIGHT versus the MAF approach for those SNPs where the source individuals are found: in Supplemental Fig S11, the overall distribution shows greater entropy per SNP for PLIGHT, while in Supplemental Fig S12, the entropy is generally higher for PLIGHT early on in the SNP removal process than for the MAF approach. This is an indicator that PLIGHT seems to be improving the degree to which a source individual can be hidden in the midst of others at an individual SNP level.

The main purpose of PLIGHT is not to determine the single, ideal method of data sanitization, but rather provide tools for users to make informed choices about the sanitization procedures that best reflect their downstream publication goals. We herein demonstrate just one interesting example of how PLIGHT may be employed. The total number of SNPs to be sanitized also depends on the data publisher’s intent for the downstream usage of the data. For example, consider a forensics study where the data publisher intends to release a sample from a known individual, but hopes to protect the individual’s relatives. The target would be to hide the source individual’s relatives (if already known or published in other databases) among background haplotypes by removing those SNPs that most strongly identify the relatives. The utility to be preserved would be the identifiability of the source individual. In the case of a functional genomics dataset, the utility to be preserved would be the genomic signal of interest, while hiding the source individual among background haplotypes (Gürsoy et al. 2020). A careful utility versus privacy analysis would have to account for the intended data usage.

We also posit a possible interactive strategy for querying datasets, where a data producer would allow an external user to query SNP genotypes from individuals in the “to-be-protected” dataset. Often the data producer would like to enable a “locus-level” query on a genome (say, by visualizing in a browser) without compromising privacy. Here, “locus-level” refers to a genomic region, potentially containing several SNPs. The question then arises as to how many SNPs can be safely released. A more basic approach is to cap the number of SNPs at a fixed amount, but this does not take into account issues such as LD or higher-order correlations between groups of SNPs, and might be too restrictive. Our approach, on the other hand, is to do the following. Based on the number and genomic positions of the SNPs, the data producer would run PLIGHT as a backend risk assessment tool to calculate the degree to which the source individual is hidden among background individuals. If the source individual is sufficiently obscured (say, is equally likely to be found as several other background individuals), the data producer can release the queried SNPs – i.e., the loci in a browser. Alternatively, the producer may prune any highly identifying SNPs and release a reduced version of the SNP genotypes to the user.

## Discussion

The analyses described have reaffirmed the prevailing notion that very few SNPs, even common ones, have the ability to leak identifying information about a query individual. Furthermore, we have shown that this leakage extends beyond the direct discovery of an individual known to be a part of a database, but can also provide piecewise genetic matches within databases, either exactly or approximately, conditional on any chosen recombination rate model. This idea of mosaic genotypic matching naturally includes the ability to identify genetic relatives within databases, and we show that our algorithm has the ability to discover 1^st^-order relatives such as parents, children and siblings (and to a lesser degree, 2^nd^-order relatives) in cohorts of size ~5,000 haplotypes with as few as 30 common SNPs on a single chromosome. The upshot of these results is that investigators seeking to release genetic or omics datasets can employ our tool to assess the degree to which their to-be-publicized data could be used to compromise the identity of a study cohort or any related individuals.

It is our contention that this risk even extends beyond the direct identification of genetic identity or relatedness. To see how, we elucidate the capacity to extract further information from inferred trajectories. The process of identifying all genotypic trajectories that match a sparse set of data can be seen as a form of coarse-grained imputation: given a few SNPs, the genomic segments containing those SNPs are extended based on available reference haplotypes. However, instead of simply identifying one best-fit genotypic extension, we provide a larger list of all genotypic extensions that are consistent with the query SNP set. In other words, we provide a sense of the “entropy” of the query genotypes conditional on a chosen reference set, where the entropy is a measure of the size of the best-fit genotypic state space in different genomic regions. An attacker could use the auxiliary information of individuals pooled across these genomic regions to explore group phenotypic risk at particular loci, for example. We presented the results for one simplified case of such an attack, but a more sophisticated usage of the pooling attack could pick out risks for both Mendelian and complex genetic disorders.

We note that previous studies have used the Li-Stephens model to carry out genetic privacy attacks, albeit by imputing SNPs at unobserved sites. For example, Samani et al (Samani et al. 2015) and Raisaro et al (Raisaro et al. 2020) both use genetic imputation based on nearby genotyped variants to find the most likely SNPs at unobserved loci. While these are powerful means of extracting further genetic information, in the case of extremely sparse sets of SNPs, the use of the conditional distributions of a set of nearby SNPs will likely not be feasible. We therefore see PLIGHT as complementary to these approaches. We have endeavored to show that, even in the absence of the ability to impute genotypes, leakage of information is possible. In fact, we foresee ways in which PLIGHT could fit into comprehensive privacy risk assessment platforms such as GenoShare (Raisaro et al. 2020) by adding complementary quantifications for sparse SNP datasets.

The expansion of genetic reference databases has also highlighted the limitations of the current reliance on single, linear reference genomes. Capturing the full range of genetic variation across a species (including SNPs, indels, structural variants (SVs), tandem repeats, etc.) in a manner that avoids biases associated with the choice of reference, requires reference data structures that are able to represent all possible sequence paths through available database genomes. Graph genomes (Novak et al. 2017; Paten et al. 2017) have been recently proposed as such frameworks for the generation of more inclusive reference databases, accounting for all known variants within a single traversable data structure. In one example (Rakocevic et al. 2019), sequences are represented as edges and breakpoints between variants as nodes. The number of branches extending out or into any node depends on the number of variants associated with that genetic locus. A single haplotype is constructed by starting at one end and traversing (in an acyclic manner) all branches that agree with the variants in the observed haplotype. If we now pose our query-matching problem in the context of graph genomes, we see that: (a) locating the individual query genotypes within the matching graph edges is easy, although we would have to include alternate branches weighted by the probability of de novo mutation or genotyping error; (b) identifying the consistent genotypic trajectories would amount to finding all possible paths that run through the branches determined in (a); and (c) it would be possible to account for linkage disequilibrium and the relative likelihood of each trajectory if the nodes of the graph genome were annotated with recombination rates calculated based on the reference database. In this way, we would be able to carry over the HMM approach into a graph genome context. If information on the number and identity of the reference haplotypes that map to each branch were also available, the pooling inference described above would also be possible.

Finally, we are also exploring the possibilities for using the underlying model for other problems. For example, the mapping of somatic mutations in cells to cancer lineages or viral mutations to clades. One interesting analysis would involve the iterative use of PLIGHT to parse through contributing genomes in a contaminated sample, by finding the most likely sets of SNPs arising from a single source genome. Ultimately, the generality and simplicity of the models make them attractive for multiple applications, as is clear from the ubiquity of HMM-based genomic analyses in the literature.

Our intention is to provide a tool to assess the degree of leakage of a given set of SNPs, and to thereby determine the privacy leakage risk associated with a dataset. This intention is, of course, complementary to subsequent sanitization procedures, such as those outlined in our previous publication (Gürsoy et al. 2020). In constructing *PLIGHT_SanitizeGenotypes*, we have envisioned one way of coupling the information from PLIGHT with data sanitization methods such as pBAM (Gürsoy et al. 2020) generation. By applying such an approach, we believe that a balance can be struck between the utility of the released dataset and the associated privacy risk.

## Methods

### PLIGHT Framework: Li-Stephens model and associated biological parameters

Let 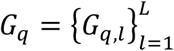 be the genotypes of a query individual *q*, observed at SNP loci *l* = {1,2, …, *L*}. The probability of observing such an individual given a space of reference haplotypes 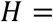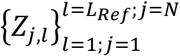 (*L*_*Ref*_ = number of genotyped sites in the reference genomes, *N* = number of haplotypes in the reference database) is:

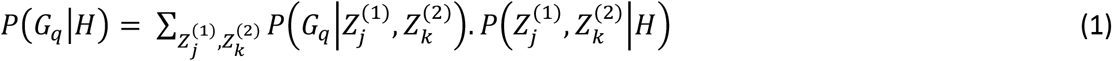

where the set of all possible haplotypes at the observed loci on the two chromosomes is given by 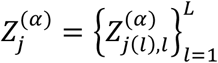, with 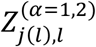 being the haplotype at position *l*, and *j* being the index of the sampled haplotype. The locus position is defined with respect to a linear reference genome, with the particular genome build assumed to be the same between the query and reference database genomes. We treat the haplotype index *j*(*l*) as a function of *l*, as it is possible for the choice of reference haplotype to be different at each locus; that is, in the haplotype matching process, recombination between reference haplotypes may occur from one observed locus to the next (see **Fig. S1**). The second subscript explicitly indicates that, from reference haplotype *j*(*l*), we select the genotype at locus *l*.

As previously described, we define a *trajectory* for a diploid genome query as 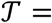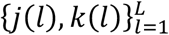, which is the sequence of reference haplotype pairs that best match the query SNPs. In the case of a haploid query, the trajectory would be a sequence of single labels at each locus.

The assumption in the current iteration of the algorithm is that the observed genotypes and the reference haplotypes are registered with respect to the same, linear reference genome. This enables a simpler matching of reference haplotypes to observed genotypes. For data structures such as personal genomes and graph genomes additional genotype matching strategies would need to be incorporated, but the conceptual framework of searching through recombining haplotypes would be the same. In general, the set of genotyped sites does not have to perfectly overlap with the set of reference haplotype sites because of rare SNPs in an individual’s genotype or differences in genotyping arrays. However, for the purposes of this study we only consider genotyped sites that overlap with those of the reference haplotypes, especially given our interest in determining the identification power of common SNPs. In case of the presence of structural variants overlapping SNP loci, we allow for missing genotypes in any of the reference haplotypes (except for *PLIGHT_Truncated* due to methodological conflicts by including missing genotypes). We thus consider the reference haplotypes as providing the complete search space, especially in light of the constantly growing genetic databases available for comparison. We avoid making explicit assumptions of population membership and statistics for the query individual with the belief that, beyond the implicit assumptions of the chosen reference set, this will enable more unbiased estimates of kinship and genotypic similarity.

The two terms on the right-hand side of Equation (1) are modeled as follows:

1. 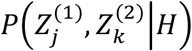 is the probability of obtaining a given set of haplotype observations at all the query loci (Li and Stephens 2003; Marchini et al. 2007):

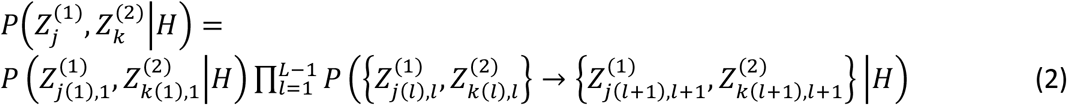
2. All terms are written with the conditional dependence on the set of reference haplotypes, *H*, made explicit. 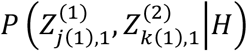 is the probability of observing a given set of haplotypes at the first query locus. This is often drawn from a uniform distribution across all haplotype pairs, but could be modified if prior knowledge on the membership of the query individual in a particular subpopulation is available. Recombination is incorporated into the analysis in the expressions for the transition probabilities from one query site to the next, 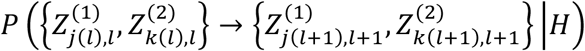 The specific expressions for the transition probabilities are provided in the Supplemental Materials. In our methods, a linear model of recombination (as in the Li-Stephens model) is included as the default. However, we allow for the inclusion of a user-defined model of recombination in its place. For example, if there is a known recombination hotspot between two adjacent query sites, it would alter the probability of transitioning between reference haplotypes in the search space, and thus impact the best-fit haplotypes calculated by the method. The user can explicitly include a vector of recombination values to be used for the *L* − 1 intervals between query sites. Additionally, we make an implicit assumption of uniform transition probabilities, where transitions between any pair of haplotypes is the equally likely. If, on the other hand, a model is to be constructed where different subgroups have distinct recombination rates at particular locations and/or are assumed to be impacted by assortative mating then transition probabilities would be conditional based on membership in these subgroups. In this iteration of our model, we do not provide a framework of this nature, but such an update would simply require the inclusion of appropriate bias terms conditional on the memberships of the initial and final haplotypes. However, we wish to emphasize that maintaining uniform transition probabilities helps prevent biased interpretations of ethnic group membership and isolation, and allows for the broad intermixing of haplotypes known to have occurred throughout human history (Narasimhan et al. 2019).
3. 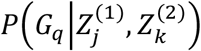 quantifies the probability of observing the query genotypes given a particular set of underlying haplotypes:

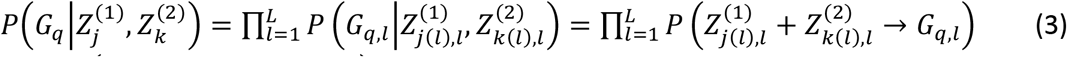

with 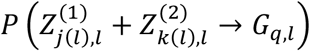 determined by the number of sites that require mutation to match the observed genotypes shown in **Table 4**. This probability helps constrain the haplotypes that are possible given the observed genotypes, allowing for the case where mutations or genotyping errors occur (as considered in IMPUTE (Howie et al. 2009)). We follow the suggestion of the authors of IMPUTE to consider a background rate of base pair mutation *θ* (the ***thetamutationrate*** parameter in our code), in addition to a mutation rate per haplotype of 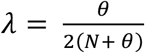 (the ***lambdamutationrate*** parameter in our code) under the assumption of a neutral coalescent tree for *N* haplotypes (Li and Stephens 2003; Marchini et al. 2007). However, it is possible for the user to explicitly augment the background rate *θ* with contributions from genotyping error, or to ignore the mutation rate altogether and set 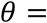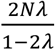 such that *λ* is equal to the known genotyping error.

**Table 4.**
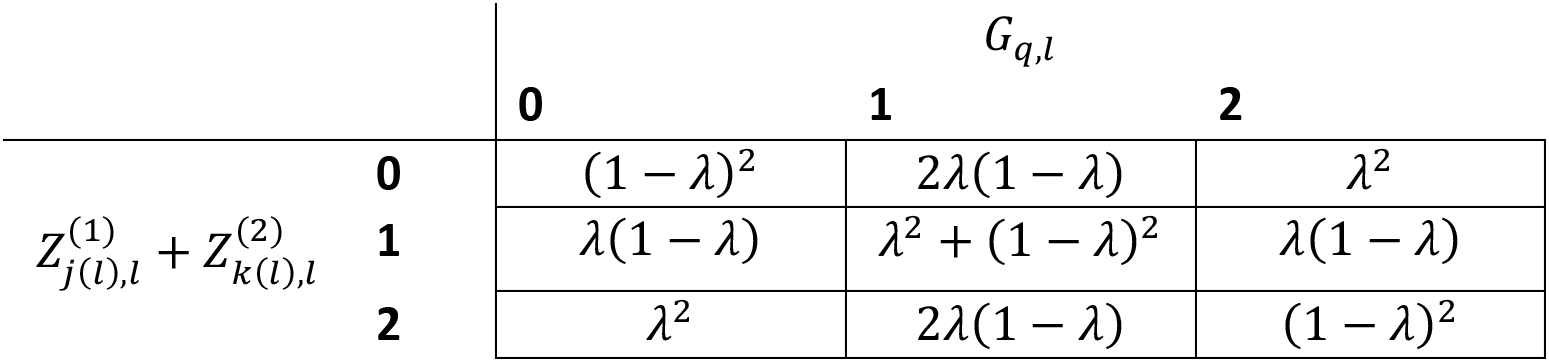
Table of conditional probabilities for the observed genotypes based on the sum of two reference haplotypes.

In summary, the aim is to figure out the contribution to the total probability of each of the haplotype combinations, by estimating 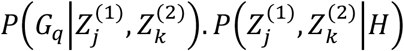 for all genotypic trajectories, and to maximize this probability.

### PLIGHT framework: Hidden Markov Model optimization

The problem of identifying the best-fit combination of haplotypes is well-suited to the framework of Hidden Markov models (HMMs) given the traditional treatment of the genome as a linear sequence of base pairs. In this understanding, meiotic recombination between loci does not occur between distant locations of a chromosome (as may occur, hypothetically, due to consistent 3D folding of the chromosomes within the nucleus), but has a certain probability of occurring at every intermediate site between any pair of loci. Usually, the greater the distance between the loci, the higher is the probability that recombination will have occurred in an ancestor of the query genome, though the probability is not necessarily uniform across every site. As seen in the previous section, it then becomes easy to associate HMM emission probabilities at genomic sites with mutation rates and HMM transition probabilities between latent haplotypes with recombination rates. Furthermore, in the above expressions first-order Markovian behavior is assumed, and the observed output genotype is seen to depend only on the underlying haplotypes at that site alone (so-called output independence). This constrains the type of HMMs considered here, but leaves open interesting future applications where such assumptions are relaxed.

The problem of identifying the best trajectory through haplotype space can be carried out using the Viterbi algorithm(Viterbi 1967). This method solves the problem of maximizing the probability of the trajectories through the latent space in time *O*((*N* × *N*)^2^*L*), where *N* is the number of possible haploid states, i.e. the number of reference haplotypes, *N* × *N* is the corresponding number of diploid states, and *L* is the number of observed loci:

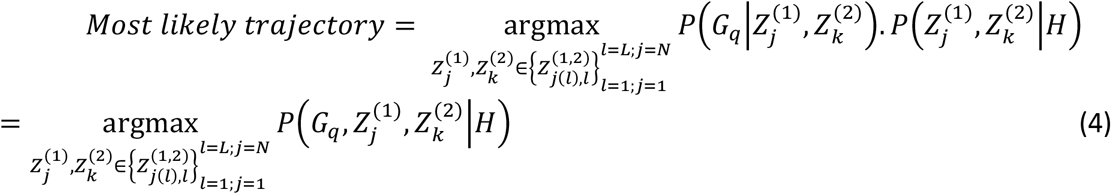

where expressions for the two probabilities are given in Equations 2 and 3. In the Supplemental Materials we provide further details on the equivalence between the optimization in Equation (4) and the Viterbi algorithm.

A calculation of the argument of maximum probability occurs separately at each pair of reference haplotypes and each locus, resulting in a set of best reference haplotypes at the previous locus. These sets of best reference haplotypes must be stored in a backtrace vector that is accessed at the last query site, when the best reference haplotypes at that site are traced back to the corresponding haplotypes from the previous site, and so on, resulting in a complete set of best-fit reference haplotypes at all observed query sites. However, the fact that the time complexity scales with the square of the number of states places strain on computational resources due to the need to account for *N* × *N* reference haplotype combinations for diploid genomes. Additionally, the memory required to store the probabilities and the backtrace states scales as *O*((*N* × *N*)*L*).

We accordingly made modifications to the Viterbi algorithm to ameliorate these pressures on computing resources:

1. We utilize the commonly employed logarithmic form of the Viterbi algorithm to prevent the accumulation and subsequent round off of vanishingly small probability products.
2. *Matrix methods.* The probability vectors were encoded as Python *numpy* arrays. Under the assumption of an unbiased transition matrix (as given in Equation S3, with no assumption of subpopulation membership nor biased recombination), each *argmax* calculation in Equation 4 was calculated over an array whose elements were updated using the following simple rules:

a. Let log *v*_*l*_(*j*, *k*) be the log-probability vector (Equation S6).
b. log *v*_*l*_(*j*, *k*) is a matrix indexed by every pair of reference haplotypes. For memory purposes this matrix was flattened in 1D, keeping only the lower triangle of the matrix.
c. At each observed genotype locus, initialize 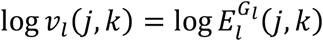, the vector of precalculated log-emission-probabilities for each pair of reference haplotypes and the observed genotype.
d. For *l* = 1, set 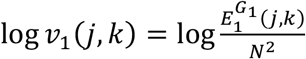, with the assumption of equal likelihood of all reference haplotypes at the first observed locus.
e. Define 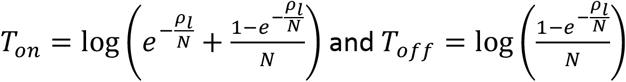
f. Define the matrix Δ(*j*, *k*) = log *v*_*l*-1_(*j*, *k*) + 2*T*_*off*_
g. For every pair of reference haplotypes (*j*, *k*), update all elements in the same row and column: Δ(*j*, .) += *T*_*on*_ − *T*_*off*_ and Δ(., *k*) += *T*_*on*_ − *T*_*off*_.
h. Find the maximum log-probability 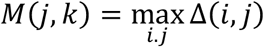 and the corresponding arguments 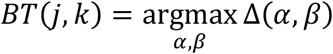, where the last term is the backtrace vector.
i. Repeat steps (f)-(h) for all haplotype pairs. Note the matrix Δ(*j*, *k*) is reinitialized every time in step (f).
j. Importantly, we modified the previous step to include all pairs of haplotypes that were within a certain range of the absolute maximum (by default, the cutoff was set as |*M*(*j*, *k*)| − 0.01 * |*M*(*j*, *k*)|. We did this as, given the assumed sparsity of the data, it was likely that multiple haplotypes would match exactly or nearly so. Additionally, this looser definition of maximization also compensates any rounding-off errors that may have caused two similar paths to deviate in log-probability. Changing this parameter could also allow the user to discover sub-optimal paths as well, if desired.
k. Update log *v*_*l*_(*j*, *k*) += *M*(*j*, *k*).
l. Repeat steps (c)-(k) for every observed genotyped site.
m. When *l* = *L*, the process is terminated by finding 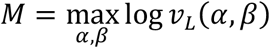 and *BT* = *All* (*j*, *k*) *such that* |log *v*_*L*_(*j*, *k*)| ≥ |*M*| − 0.01 * |*M*|.
n. Using this terminal set of *BT*, the corresponding backtrace values for each selected pair of reference haplotypes are chosen and traced all the way back to the first observed site. This results in a set of trajectories that may fork and merge, and which form the basis of the subsequent phenotypic inference.
3. *Memory constraints.* The scaling of the matrices with the number of pairs of reference haplotypes puts significant constraints on the reference database that can be considered at any point in the analysis. We discuss two of the modules constructed to address memory issues in later subsections, and focus here on the treatment of the backtrace vector used in all the modules. Instead of explicitly storing the backtrace vector in RAM, we use the Python package *gzip* to write the backtrace vector determined for each observed site directly to a gzipped file. We used the same strategy to read through the gzipped file during the final stage of recovering the best-fit trajectories.

### PLIGHT Framework: Parallelization, Truncation and Iterative Schemes

We created 3 separate modules in our package: first, the module *PLIGHT_Exact* encodes the full Li-Stephens HMM as described above; second, we created a module *PLIGHT_Truncated* that truncates the possible trajectory extensions at each observed site to a fraction of the total possible state space to reduce the memory requirements; third, we created a module *PLIGHT_Iterative* that slices up the total reference space into randomly chosen, more tractable subsets, runs *PLIGHT_Exact* on the subsets and pools the best states for a rerun. *PLIGHT_Truncated* was created to consider means of reducing the memory footprint on hard drives, as the backtrace vectors in the HMM model can occupy a significant amount of space. However, the amount of RAM consumed by *PLIGHT_Truncated* remains essentially the same as *PLIGHT_Exact*. The *PLIGHT_Iterative* algorithm was designed to ameliorate both the hard disk and RAM costs of the model, and is thus the preferred method for reference databases beyond ~500 reference haplotypes. In the following, we describe both the parallelization procedure for speed-up, and the additional steps taken in the last two modules, which are, in general, approximations of the full model. However, we show that the approximations often approach the full model when judicious choices of the parameters are made. Additionally, for the first module, we explicitly design a function for the case where the genotyped individual is known to lie within the reference database, as this problem involves a significantly more restricted search (being one-dimensional in the search for genotypes versus the two-dimensional case of pairs of haplotypes), with the recombination rate set to 0. This version of the *PLIGHT_Exact* algorithm is referred to as the *PLIGHT_InRef* algorithm.

#### Parallelization scheme

In all three modules, we enable the usage of Python’s *multiprocessing* scheme to run the calculations over every set of haplotype pairs in parallel. This greatly improves the speed of the analysis, as the matrix manipulations are some of the more time-consuming steps.

#### Truncation scheme

In the *PLIGHT_Truncated* module, at every observed site we truncate the possible haplotype states by choosing the top *T* sets of (*α*, *β*) pairs. This scheme was inspired by similar techniques employed in the Eagle2 imputation program (Loh et al. 2016). The main premise of this approximation is after a certain number of observed loci, only a fraction of a the total number of trajectories will meaningfully contribute to the best-fit states, and allow the retention of only a fraction of the total number of states in memory. We discuss the details of this method and its associated results in the Supplemental Materials, as it mainly serves to demonstrate the compressibility of many trajectories.

#### Iterative scheme

In the *PLIGHT_Iterative* module, the following sequence of steps are run:

a. Set a tractable subgroup size *S*_*sg*_ for a single run of the full HMM model
b. Randomly shuffle the identities of the reference haplotypes, and chunk the full reference haplotype into subgroups of size *S*_*sg*_.
c. Run the full HMM model as described in the section *Hidden Markov Model optimization* for each subgroup.
d. Repeat steps (b) and (c) a user-defined *n*_*iter*_ times.
e. Pool together all the best-fit haplotypes from each of the subgroups and each of the iterations; best-fit pairs are separated into their constituent haplotypes at this stage.
f. This pooled set is fed back into step (b) and the process is repeated until:

i. The length of the pooled list is smaller than *S*_*sg*_; or
ii. The current pooled list is identical to the list derived during the previous pooling step; or
iii. The current pooled list is larger in size than the previous pooled list.
g. Once the outermost loop over pooling steps is exited, the full module is run on the final best-fit list of haplotypes, with the output being a file with the best-fit pairs of haplotypes chosen from the final pooled list.
h. The parallelization is run over the calculations in step (c) as before.

The trade-off here is, of course, between speed and memory usage. Searching through large databases could take significantly longer even when distributed in this manner, but the memory burden is substantially alleviated.

An issue related to the subdivision process is that the globally optimal combinations of haplotypes, as obtainable in an exact Li-Stephens HMM algorithm, may not co-occur in the subgroups defined in this approximate algorithm. This “mixing problem” is dealt with in two ways: for the inner loop, we run the (subdivision + Exact HMM) process *n*_*iter*_ times, with each iteration involving a random subdivision of the input haplotypes and a subsequent running of the HMM for each subdivision in each iteration (total HMMs run: *n*_*iter*_ × *n*_*subgroups*_, with *n*_*groups*_ = number of subgroups required to include all input haplotypes at this stage); for the outer loop, we obtain the union of the best-fit haplotypes across all *n*_*iter*_ × *n*_*subgroups*_ runs and use this set of haplotypes as the input set for the next round of random subdivisions. Note that for the union set in the outer loop, we only include the identities of the haplotypes and none of the haplotype combinations from the inner loop HMM runs. The idea is that if certain haplotypes have significant contributions to any segments of the genotype match, they will be retained through the different stages and allowed to combine with many other haplotypes. The loops are exited if there is no change in the set of best-fit haplotypes, or if the best-fit list from one iteration of the outer loop is larger than that from the previous iteration (to prevent infinite loops of iteration), or if the set of best-fit haplotypes can be determined by a single run of *PLIGHT_Exact*. If the loop exits with a large number of best-fit haplotypes, we recommend rerunning the code (to allow the randomness to explore different trajectories) or modifying the parameters (such as the recombination rates or *n*_*iter*_).

### PLIGHT Framework: Visualization and analysis module

To help visualize the full set of trajectories through the reference haplotype space, we constructed a processing and visualization module, termed *PLIGHT_Vis*. For the trajectory representation step, the most efficient way of representing the multiple possible trajectories with potential overlap was to create a series of multiply-linked lists. This involved reading through the output list of best-fit nodes from the HMM modules, with each set of nodes being laid down in the graph in reverse order from the final observed genotype site to the first. At every observed site, the union set of nodes (i.e. with no repeated nodes, even if the same node appears in several parallel trajectories) are laid out and their connectivity with the soon-to-be-added layer at the previous site is established. This continues until the whole graph is constructed. The Python package *matplotlib* is then used to generate a visual representation of the graph with the help of the linked lists. Arrows connect nodes (represented as dots) at one observed site to their corresponding antecedents at the next observed site. The reference sample identities of the two inferred haplotypes at each node are printed above and below the dots.

Additionally, we provide two simple analysis tools as well:

1. For each chromosome, we identify the maximally represented inferred haplotype at each observed site and provide this information at the top of the visualization.
2. For cross-chromosome quantification of the most representative trajectories, we score each trajectory in each chromosome by a weighted sum,

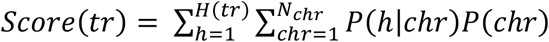

where *H*(*tr*) = Number of unique haplotypes within trajectory *tr*, *P*(*h*|*chr*) = Probability of instances of haplotype *i* occurring in the predicted trajectory set for chromosome *chr*, *P*(*chr*) = Probability of each chromosome, taken to be a uniform distribution 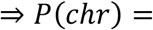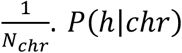 is calculated as the fraction of trajectories for a given chromosome within which the haplotype *h* occurs. The score of each trajectory is therefore the sum of the cross-chromosome probability of a haplotype being found in the prediction, taken over all unique haplotypes in that trajectory. The cross-chromosome *consensus* trajectories with the maximum score in each chromosome are stored in a file. This weighting heuristic was chosen with the intention of identifying trajectories in each chromosome that share significant information with trajectories in all other chromosomes.

### PLIGHT Framework: Quantitative Metrics

We output four quantitative metrics for each of the algorithms. The first is the log-probability associated with the most likely trajectories, 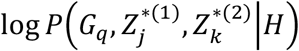, where 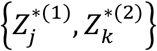 represent the most likely trajectories. This metric is dependent on the number and MAF of the SNPs, as well as the degree of matching to the reference database haplotypes. Accordingly, this metric is not comparable across query SNP sets, but can be used to diagnose whether the same query set matches differently to a different reference database or with a different run of *PLIGHT_Iterative* (which has a certain degree of stochasticity).

The second metric is the logarithm of the joint likelihood of the SNPs conditional on the HMM model under consideration:

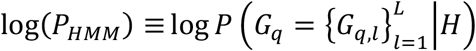

This value is calculated as the total probability of the last step of the HMM,

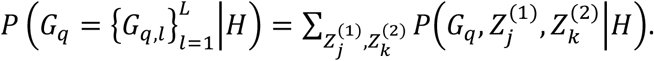

The third metric is the logarithm of the likelihood of observing the SNPs assuming they are all under Hardy-Weinberg equilibrium (HWE):

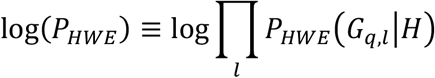

where 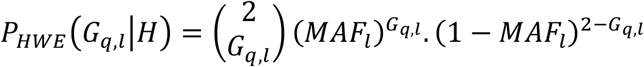

The final metric is the logarithm of the likelihood of observing the genotypes assuming independence, i.e. instead of the alleles being under HWE, this just assumes that reference database captures the true independent probabilities of the diploid genotypes:

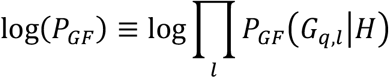

where *P*_*GF*_(*G*_*q,l*_|*H*) = *Frequency of occurrence of G*_*q,l*_ *in the reference population* The log(*P*_*GF*_) is especially relevant for *PLIGHT_InRef*, where the genotypes are matched and not pairs of haplotypes.

### PLIGHT Framework: Sanitization Module

To help guide the decisions behind which SNPs to publish, we provide a simple tool for pruning a single SNP from the query set, based on the results of running PLIGHT. This tool, termed *PLIGHT_SanitizeGenotypes*, operates as follows:

1. The best-fit trajectories from a previous PLIGHT run are passed as input.
2. The SNPs for which the number of unique haplotype pairs (*N*_*UHP*_) is the least are identified (see Supplemental Fig S8). That is, for each SNP, the total number of unique haplotype pairs in the trajectories at that locus are counted, and those SNPs which have the smallest number are flagged. These subsets of query SNPs have the lowest “entropy” in terms of reference database matches, and are likely to contribute to easy reidentification. Formally, *N*_*UHP*_(*l*) = *No. of unique pairs of* {*j*(*l*), *k*(*l*)} *in the best* − *fit trajectories at locus l*.
3. We then rank these SNPs by MAF, and remove the one with the lowest MAF from the query list. A new, pruned query list is then generated and run through PLIGHT again.

This process can be repeated for every SNP a user chooses to remove.

We describe the particular strategies employed for the demonstration in the *Results* section. The metrics considered are defined as follows.

1. Total informational entropy of the identified individuals: 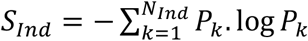, where *N*_*Ind*_ = *No*. *of individuals across all trajectories in a single inference run*, and *P*_S_ = *Probability of observing individual k across all trajectories in a single inference run* This is a measure of the diversity of individuals identified in an inference run.
2. Maximum probability of observing any of the two source individuals in the trajectory: 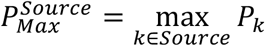, where *P*_S_ is defined in the preceding. This is a measure of how identifiable (either of) the source individuals are.
3. Per-SNP entropy of identified individuals, if either of the source individuals is found at that 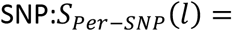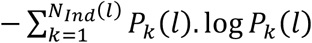. log *P*_S_(*l*) *if the source individual is found at SNP i*, where *N*_*Ind*_(*i*) = *No*. *of individuals at SNP i*, and *P*_S_(*i*) = *Probability of observing individual k across all trajectories at SNP i*. This is a measure of the observed diversity in inferred individuals at each SNP position.

### Simulation of samples

Our evaluation of the performance of the code was often conducted on simulated data. For the analysis of individuals known to be within the reference database, we selected a single individual at random from the reference database. For the analysis of mosaic genotypes, we selected two or more individuals at random. For all the individuals in a given simulation we created a genotype sample using the following methods. To spread out the selected SNPs across the genome, we only choose SNPs ordered along the chromosome with a probability *p*~*Bernoulli*(1,0.003). Each SNP thus chosen is randomly mutated at the rate 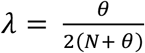 at each site (i.e. each SNP has a probability of being mutated to a new value given by Table 4). If the final genotype at the SNP is heterozygous or homozygous in the alternate allele, we select that SNP in our final list. Note that this selection process is meant to mimic the case where only alternative alleles are obtained, and the reference alleles are left out. However, this can also lead to an apparent inflation of the mutation rate, as unmutated, homozygous reference alleles are left out. In the results shown below, this inflation of the apparent mutation rate is implicitly assumed.

For the individual-in-the-reference case, this sampling method directly provides the input dataset. For the mosaic case, we divide each chromosome into a set of segments (for example, two segments for a mosaic of two reference individuals) and assign the genotypes of different individuals, mutated in the above fashion, to each of the segments.

### Mosaic genome reconstruction

To validate the inferred trajectories against the query genomes, we need to reconstruct the mosaic genomes based on the labels inferred. To do so, we essentially grab segments from the reference vcf file corresponding to the reference haplotypes inferred in each trajectory: for 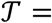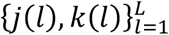, we have *L* − 1 segments to reconstruct; for each segment (defined in the following), we extract all the SNP haplotypes for reference haplotypes *j* and *k*, combining their values at each position to get the inferred genotype. The genomic segments of identified individuals at each SNP were constructed according to: *Start of segment* 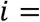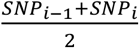; *End of segment* 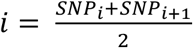; where *SNP*_*i*_ = *Genomic position of SNP i*, and the first segment started at *SNP*_1_ and the last segment ended at *SNP*_*L*_. We store the results of this reconstruction in a vcf file.

### Analysis of SNPs derived from coffee cups

The query SNPs derived from a single genotyped individual were obtained from samples collected as part of our previously published analysis (Gürsoy et al. 2020). These include Illumina Whole Genome Sequencing (WGS) results from swabs of coffee cups, as well as gold-standard blood tissue samples from the same individuals. In brief, a QIAmp DNA Investigator Kit was used to purify DNA from the coffee cup swabs; the DNA was PCR-amplified using a REPLI-g Single Cell Kit; for the tissue samples, purified PCR-free DNA was directly used; all samples subsequently underwent Illumina WGS. The resulting fastq files were mapped to the hg19 reference genome (b37 assembly) using bwa (Li and Durbin 2009), with the BAM outputs being de-duplicated using Picard tools. Finally, the de-duplicated BAM files were processed utilizing GATK (DePristo et al. 2011; Van der Auwera et al. 2013) to produce variant call sets in vcf format from both genotyping assays. For further details on sample and data processing, see ref. (Gürsoy et al. 2020). All genotypes were unphased, and so when the blood tissue sample is used as a reference, we arbitrarily phase the data; e.g., a genotype call of “0/1” gets distributed with “0” to the first haplotype and “1” to the second.

The coffee-cup-based genotypes serve as the noisy, sparse dataset, while the blood-tissue-based results form the higher quality baseline for comparison. The vcf coordinates are defined according human genome reference assembly *GRCh37*, using the reference fasta file *human_g1k_v37.fasta.gz (*ftp.1000genomes.ebi.ac.uk/vol1/ftp/technical/reference), the same as for the 1000 Genomes Phase 3 database. This registers the vcf files from the coffee cups and from the reference database of the 1000 Genomes project in the same coordinate system, an essential requirement for *PLIGHT* to identify matches between the query and reference datasets. Using mosaic genome reconstructions, we assessed the accuracy to which we could impute all SNPs across the full range of observed SNP loci. The accuracy metrics included a straightforward calculation of the fraction of SNPs correctly identified (only exact matches of the genotypes), as well as measure of the degree to which the inferred trajectory matched the query genome, with the contribution from each SNP weighted by a function of the genotype frequency:

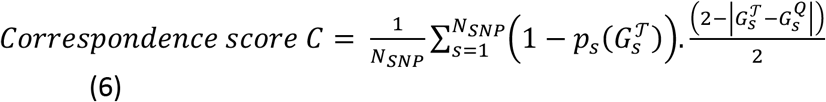

The score *C* finds the total degree of correspondence between the set of query individual genotypes, 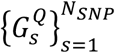, and the set of genotypes for trajectory 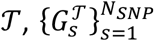, where *N*_*SNP*_ is the total number of overlapping SNPs defined in the vcf files of the reference database and the query individual between the first observed SNP and the last observed SNP. Next, 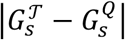 quantifies the deviation of the genotype dosage of 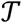 from that of the query *Q* at SNP position *s*, and is subtracted from 2 and divided by 2 to set a score scale where 0 corresponds to maximal deviation of 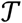 from *Q* and 1 corresponds to a perfect match between the two. Finally, 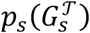 is the genotype frequency (as opposed to the allele frequency) of the SNP dosage 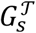, which is the probability that trajectory 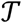 could have a given dosage at random based on population occurrence frequencies. The heuristic 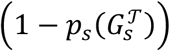 is therefore a measure of the non-randomness of the trajectory SNP dosage. In summation, *C* = 0 would occur if no SNPs matched between 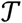 and *Q* and/or the SNPs occurred in the reference population at 100% frequency, while *C* = 1 would occur if 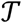 and *Q* agreed at every SNP position and the SNPs were extremely rare (and so the matching of the two is very likely to be a non-random occurrence).

We compared the fraction of correct SNPs and the correspondence scores for our trajectories to the equivalent scores calculated on a set of 99 randomly selected genomes from the 1000 Genomes database to assess if these scores were significant.

### Polygenic risk score calculation

To quantify some of the risks associated with the pooling of information across multiple genotypic trajectories, we performed approximate calculations of the linear polygenic risk scores (PRSs) based on all the SNP associations in the GWAS catalog version 1.0.2 (Buniello et al. 2019). We first identified all the individuals in each of the trajectories of a single HMM run. We then constructed the diploid mosaic genome of the query individual based on each HMM trajectory as described in the previous subsection on mosaic genome reconstruction. The resulting genotypes are then used to calculate the PRSs for each phenotype *y* and each individual *n* as:

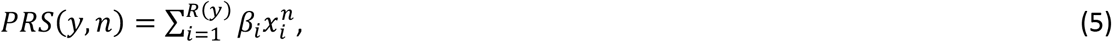

where *β*_*i*_ = *Signed effect size of the risk allele at SNP i*,
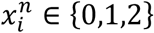 = *Genotype of individual n at SNP i*,
and *R*(*y*) = *Total number of risk alleles for phenotype y*.

The PRS is very approximate in the sense that no SNP filtering was conducted beyond those presented in the GWAS catalog, either by p-value or by LD with other associated SNPs. However, the aim here is merely to determine whether there are aggregate properties across the genome that can be inferred using our approach. We calculated the Pearson’s correlation between the PRSs of the true samples and the best-fit mosaic genomes within the regions and chromosomes sampled. All traits for which the PRS of the true sample was non-zero (‘non-zero traits’) were included. To assess whether the PRS correlations between the true individual and inferred mosaic genomes were statistically significant, we sampled a background set of ~100 individuals from the 1000 Genomes dataset that did not occur in any of the test sets we ran, and calculated the PRSs for the non-zero traits.

We ran several statistical tests to assess the correspondence in PRSs between the true samples and the best-fit mosaic genomes, relative to the background scores: (1) evaluated the cosine similarity between the true and mean values of the best-fit scores, and compared to the mean value of the background scores; (2) carried out the same analysis as in (1) but only for those traits that had more than one GWAS SNP in the regions sampled (so as to remove traits that trivially had a single SNP); (3) carried out the same analysis as in (1) but only for those traits for which the true sample had an absolute Z-score > 2 (i.e. traits for which the true sample itself is an outlier relative to the background); and (4) evaluated the cosine similarity between the true sample and each of the best-fit mosaic genomes, and then found the mean of the cosine similarity values (same for the background).

For the “Height” phenotype PRS analysis, we engineered query sets for chromosomes 3 and 6 by identifying the “Height” GWAS SNPs on the respective chromosomes as represented in the GWAS catalog. We found all SNPs within ±2 kb of the GWAS loci, and then randomly selected 90 SNPs per chromosome as our query sets. We ran *PLIGHT_Iterative* with *n*_*iter*_ = 20 and *S*_*sg*_ = 300 using the full 1000Genomes cohort, constructed the mosaic genomes from the best-fit trajectories, and calculated the PRSs for those GWAS SNPs that were within the same ±2 kb windows of our query SNPs. The result was a PRS calculation over 26 GWAS SNPs for both chromosomes (it was coincidental that the GWAS SNP numbers were the same for both chromosomes). We ran the two-sided one- or two-sample t-tests using the base R function *t.test*.

## Data Access

We have made our software available for download at https://github.com/gersteinlab/PLIGHT. We provide information on the software requirements, parameter options and examples on how to run the code. We also provide timing benchmarks for each of the algorithms. In addition, the source code is provided as a compressed folder containing the Python scripts, uploaded as Supplemental Materials.

## Acknowledgments

The authors would like to thank Hussein Mohsen for his helpful comments on the manuscript.

