## Supplemental Materials for "Assessing and mitigating privacy risk of sparse, noisy genotypes by local alignment to haplotype databases"

### Supplemental Figures

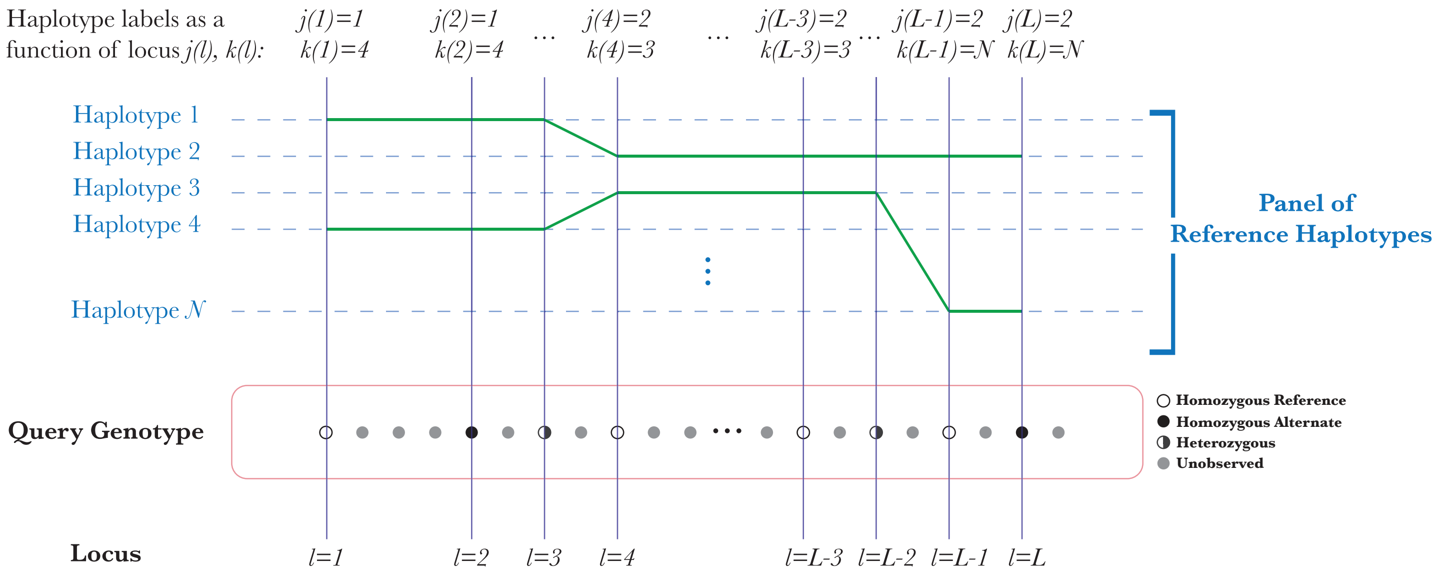


**Supplemental Figure S1.** Explanation of the j(l) indexing as a function of the query locus $l$, for one trajectory. The best-fit states in the HMM consist of a pair of haplotype labels at each locus loci $l = \{1,2,\cdots,L\}$: these are the names of the reference haplotypes that best match the query SNPs over certain genomic tracts, and the trajectory is the sequence of all such pairs of haplotype labels, $\mathcal{T} = \{j(l), k(l)\}_{l=1}^{L}$.

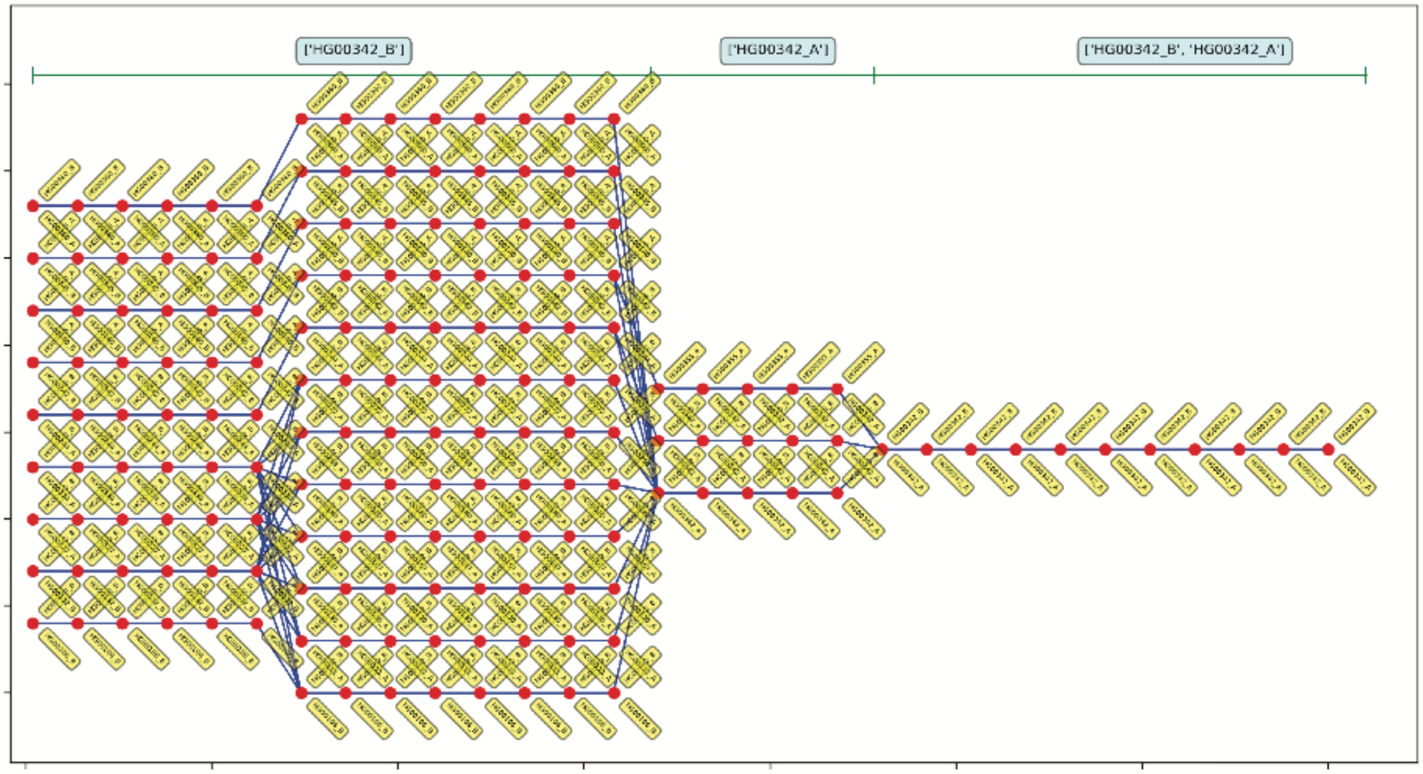


**Supplemental Figure S2.** Best-fit genotypic trajectories from PLIGHT_Exact for the diploid mosaic genome of HG00360+HG00342 constructed across 30 SNPs each for chromosome 21 (corresponding results for chromosomes 1 and 2 in Figure 3). The composition of the best-fit pair of haplotypes at each locus is depicted by two yellow tags, one below and one above the red dots.

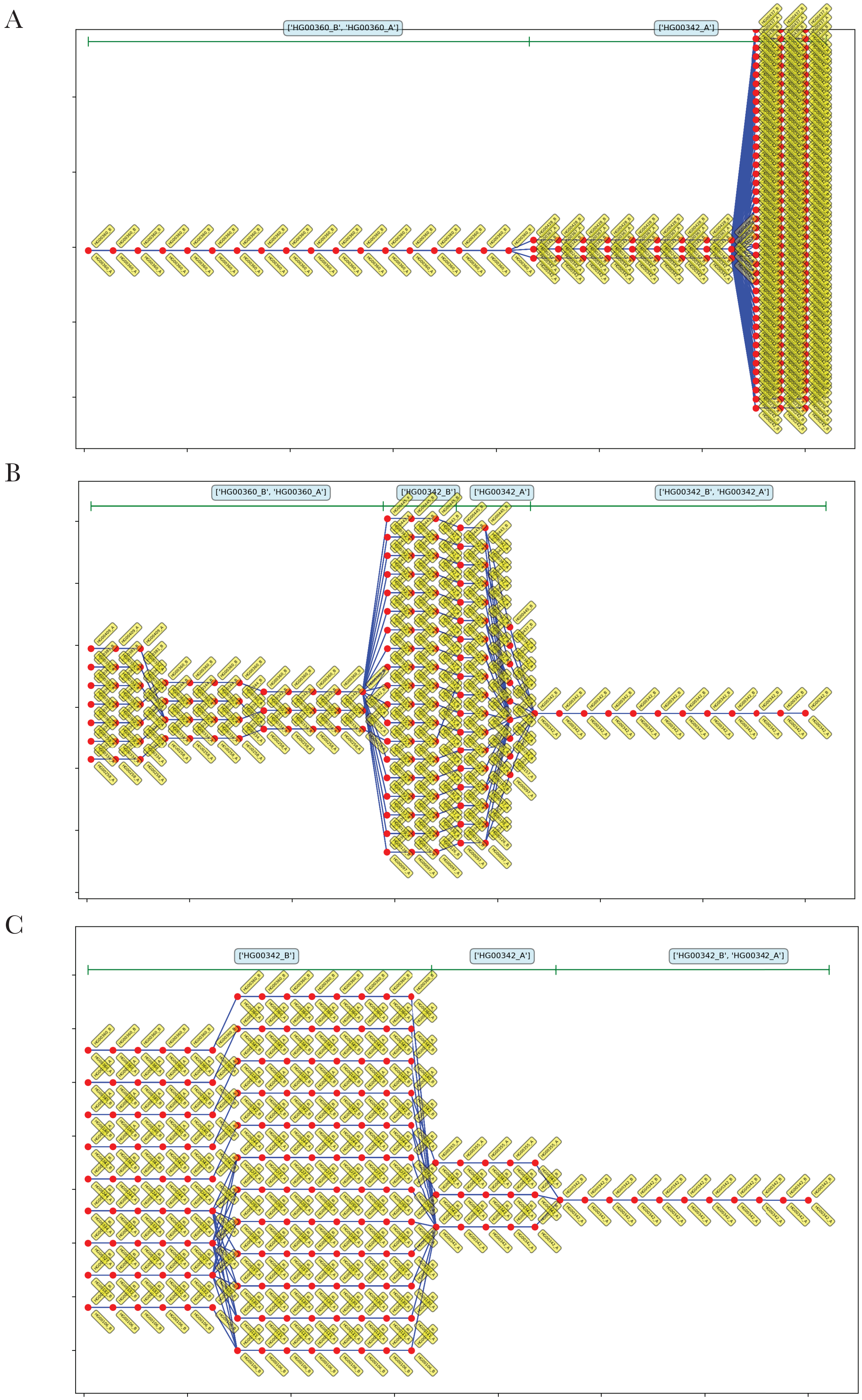
A
['HG00360_B', 'HG00360_A']
['HG00342_A']
B
['HG00360_B', 'HG00360_A']
['HG00342_B']
['HG00342_A']
['HG00342_B', 'HG00342_A']
C
['HG00342_B']
['HG00342_A']
['HG00342_B', 'HG00342_A']

**Supplemental Figure S3.** Best-fit genotypic trajectories from PLIGHT_Exact for the diploid mosaic genome of HG00360+HG00342 constructed across 30 SNPs each for chromosome with recombination rate = 1.0 cM/Mb: (A) Chromosome 1; (B) Chromosome 2; (C) Chromosome 21. The composition of the best-fit pair of haplotypes at each locus is depicted by two yellow tags, one below and one above the red dots.

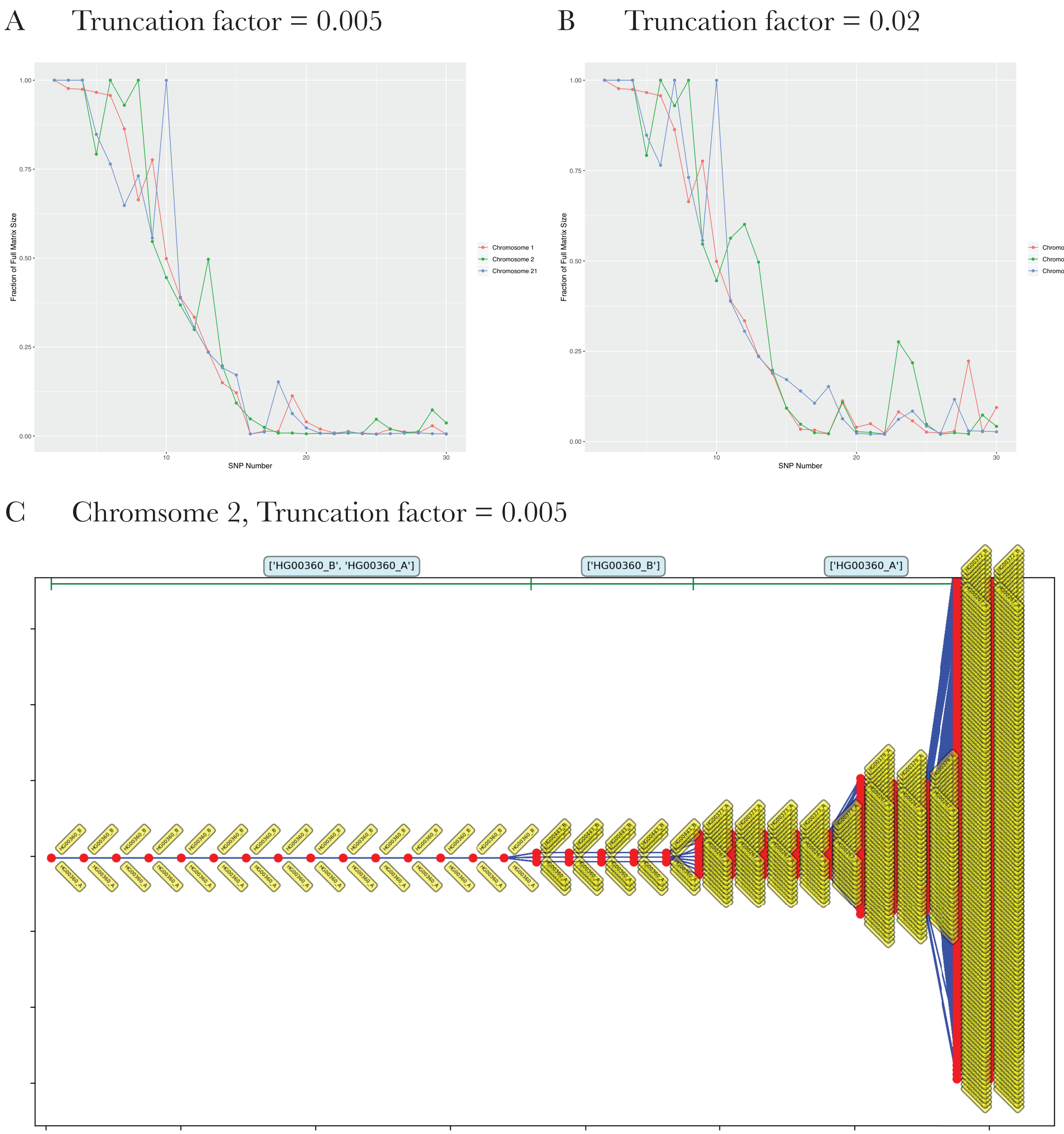


**Supplemental Figure S4.** Results for the PLIGHT_Truncated algorithm applied to the same query SNP set as for **Figure 3**. (A) Fraction of the full matrix size at each step (i.e. SNP) in the HMM sequence, for a final truncation factor of f = 0.005. (B) Fraction of the full matrix size at each step (i.e. SNP) in the HMM sequence, for a final truncation factor of f = 0.02. (C) Trajectory for chromosome 2, shown to illustrate a case where truncation results in different trajectories relative to the exact case (compare to **Figure 3B**). The composition of the best-fit pair of haplotypes at each locus is depicted by two yellow tags, one below and one above the red dots.

A

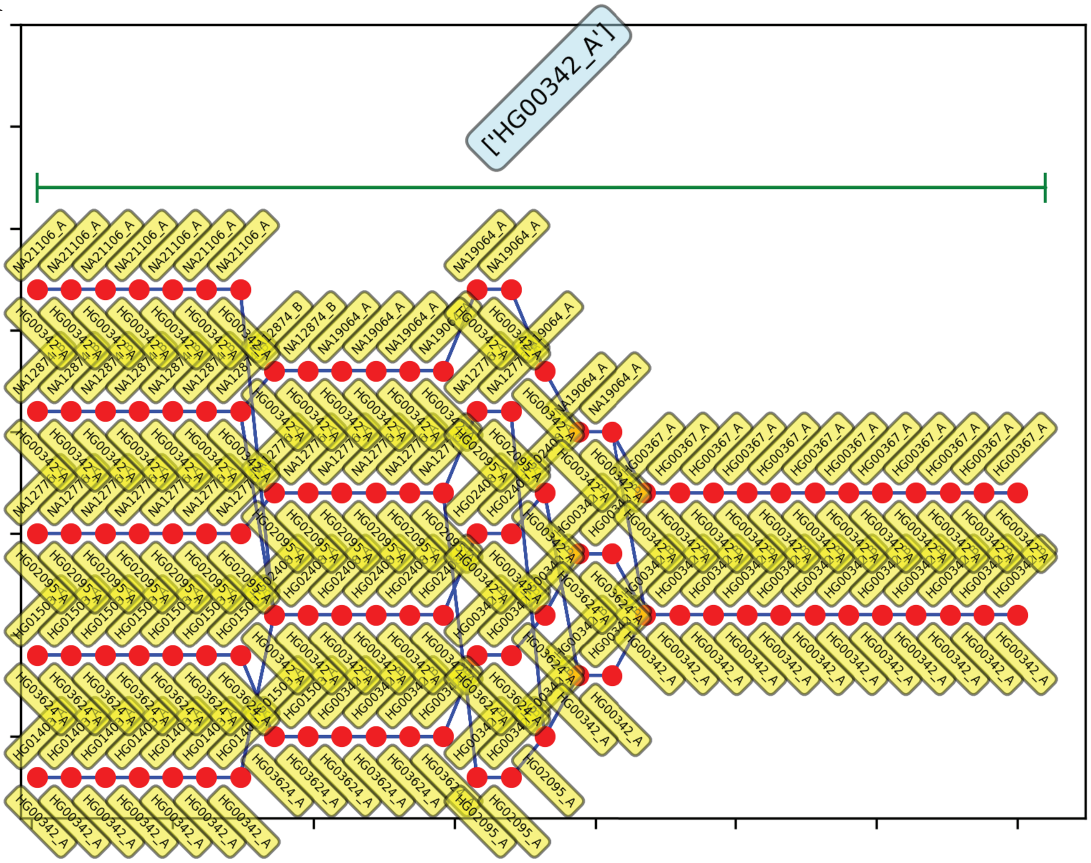


B

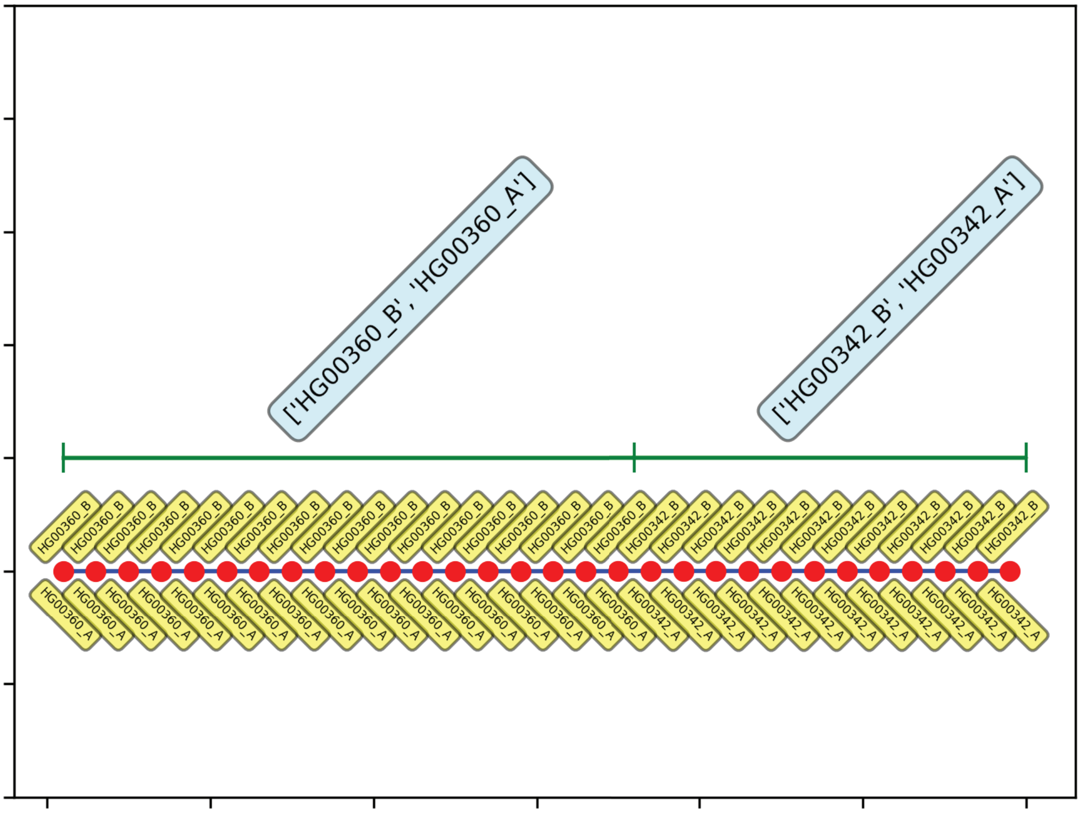

**Supplemental Figure S5.** Consensus genotypic trajectories from PLIGHT_Iterative for the diploid mosaic genome of HG00360+HG00342 constructed across 30 SNPs in chromosome 1, where the consensus score is evaluated by weighting the haplotypes in each trajectory in proportion to their occurrence across all three chromosomes. The composition of the best-fit pair of haplotypes at each locus is depicted by two yellow tags, one below and one above the red dots. (A) $\mathrm{n}_{\mathrm{iter}} = 20$, replicate 2; (B) $\mathrm{n}_{\mathrm{iter}} = 30$, replicate 2. Shown at the top of each panel are the most frequent haplotypes within each segment indicated. The corresponding first replicates are shown in Figure 4 of the main manuscript.

A

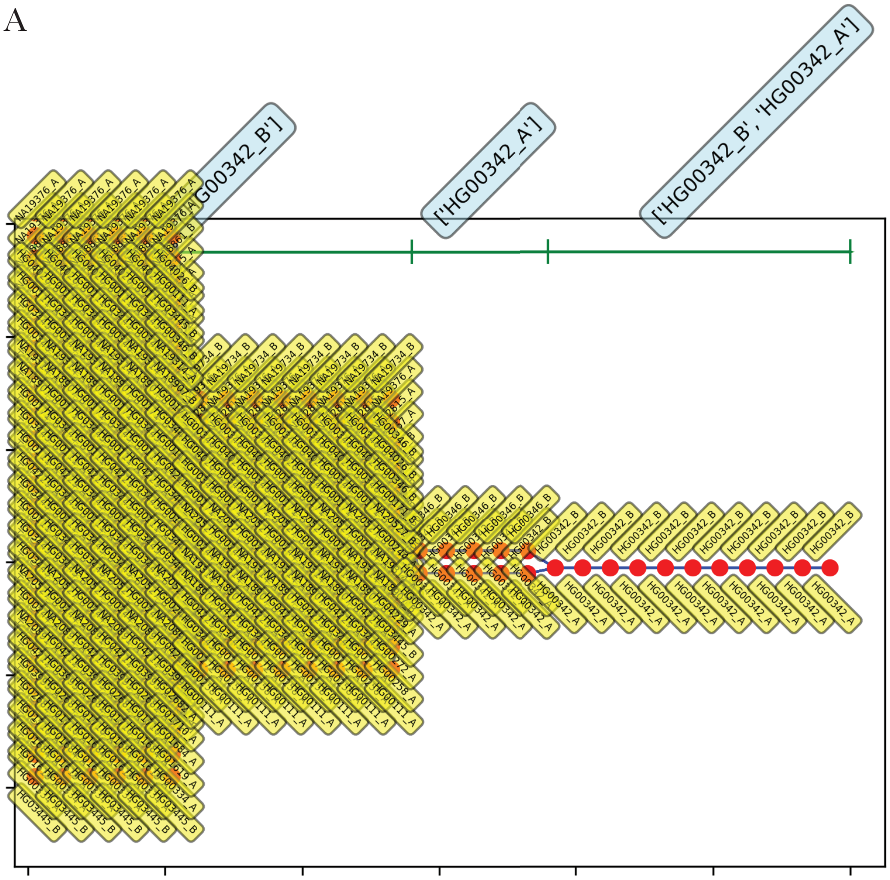
['HG00342_A']
['HG00342_B', 'HG00342_A']

B

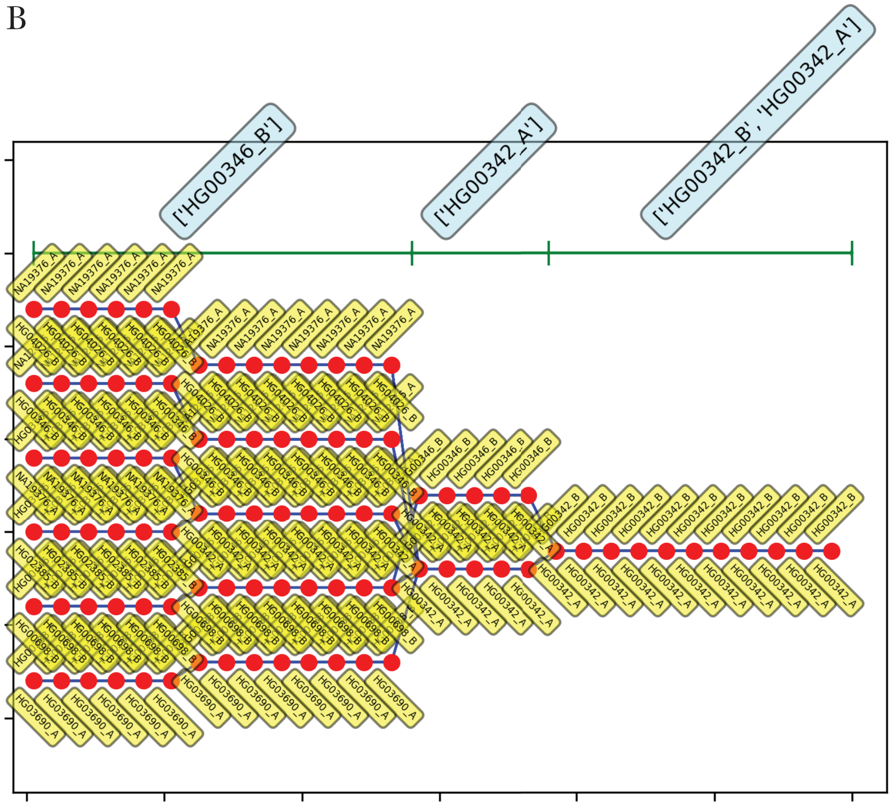
['HG00346_B']
['HG00342_A']
['HG00342_B', 'HG00342_A']

**Supplemental Figure S6.** Consensus genotypic trajectories from PLIGHT_Iterative for the diploid mosaic genome of HG00360+HG00342 constructed across 30 SNPs in chromosome 21, where the consensus score is evaluated by weighting the haplotypes in each trajectory in proportion to their occurrence across all three chromosomes. The composition of the best-fit pair of haplotypes at each locus is depicted by two yellow tags, one below and one above the red dots. (A) $\mathrm{n_{iter}} = 20$, replicate 1; (B) $\mathrm{n_{iter}} = 30$, replicate 1. Shown at the top of each panel are the most frequent haplotypes within each segment indicated. The corresponding first replicates are shown in Figure 4 of the main manuscript. Panel A is an example of a successful identification of the two component individuals, while Panel B shows a case where only one of the two individuals is found.

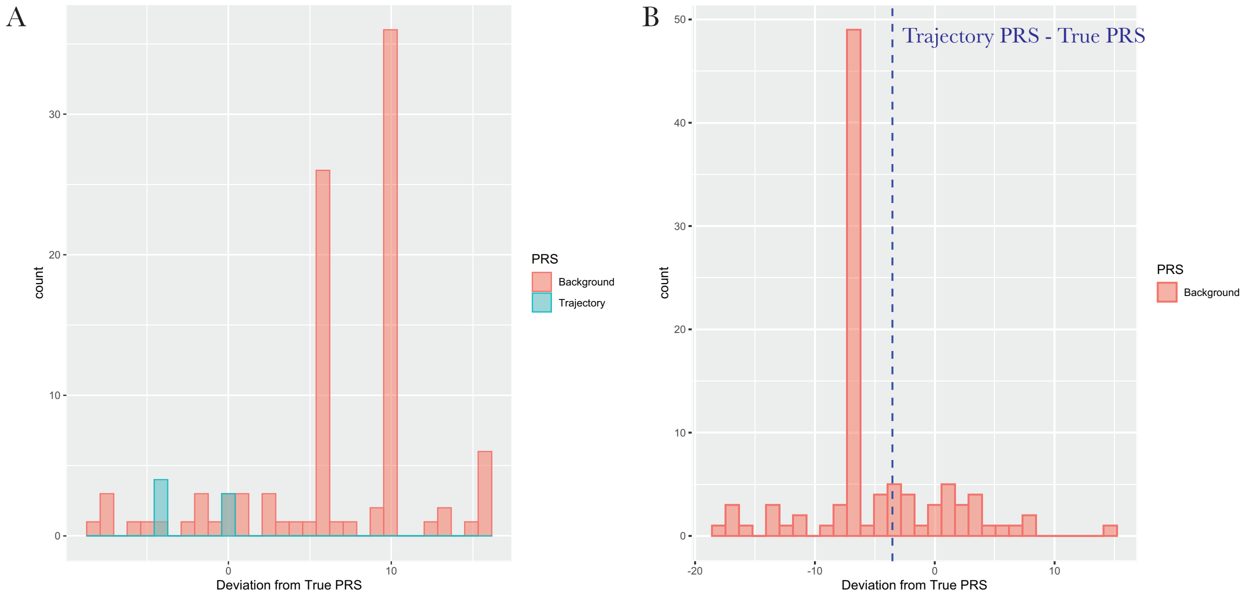


**Supplemental Figure S7.** Histograms of the deviation values from the true PRS scores for the Height GWAS analysis, shown for both inferred trajectories and background individuals. (A) Results for chromosome 3 (red = Background PRS - True PRS, blue = Trajectory PRS - True PRS); (B) Results for chromosome 6 (red = Background PRS - True PRS, blue dashed line = Trajectory PRS - True PRS).

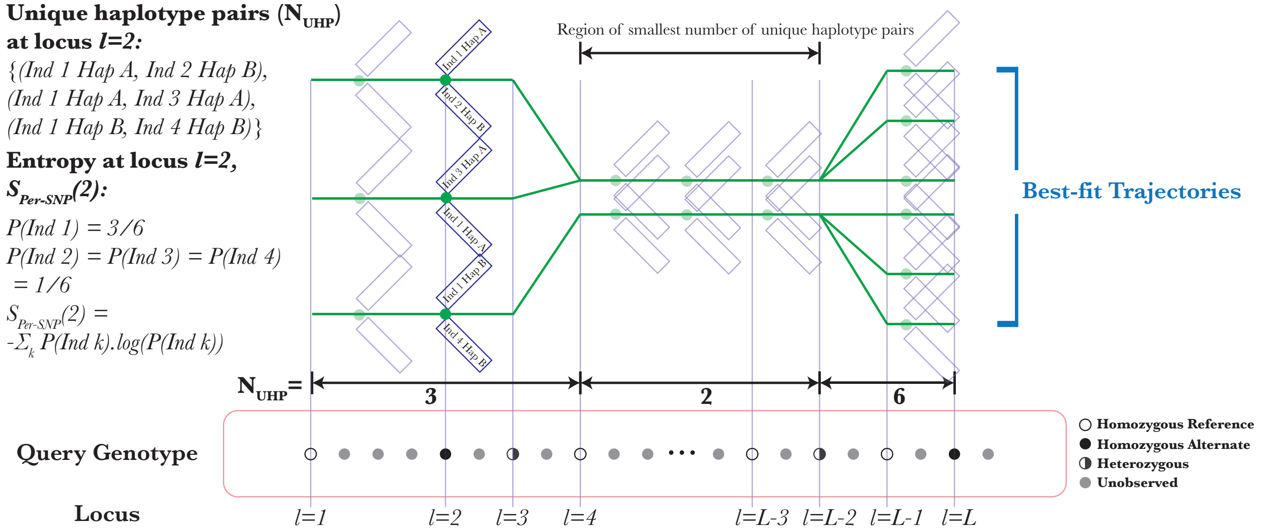


**Supplemental Figure S8.** Schematic of the metrics used in the sanitization scheme. In green is shown an example of a set of best-fit parallel trajectories. At locus l = 2, the figure shows examples of the identified haplotype pairs for this particular set of trajectories. On the left, the figure includes an explanation of how the unique haplotype pairs are counted, as wells as how the entropy across individuals per SNP is calculated at a particular locus. The figure also includes a marking of the region of the smallest number of unique haplotype pairs, which is then used in the sanitization procedure outlined in the paper.

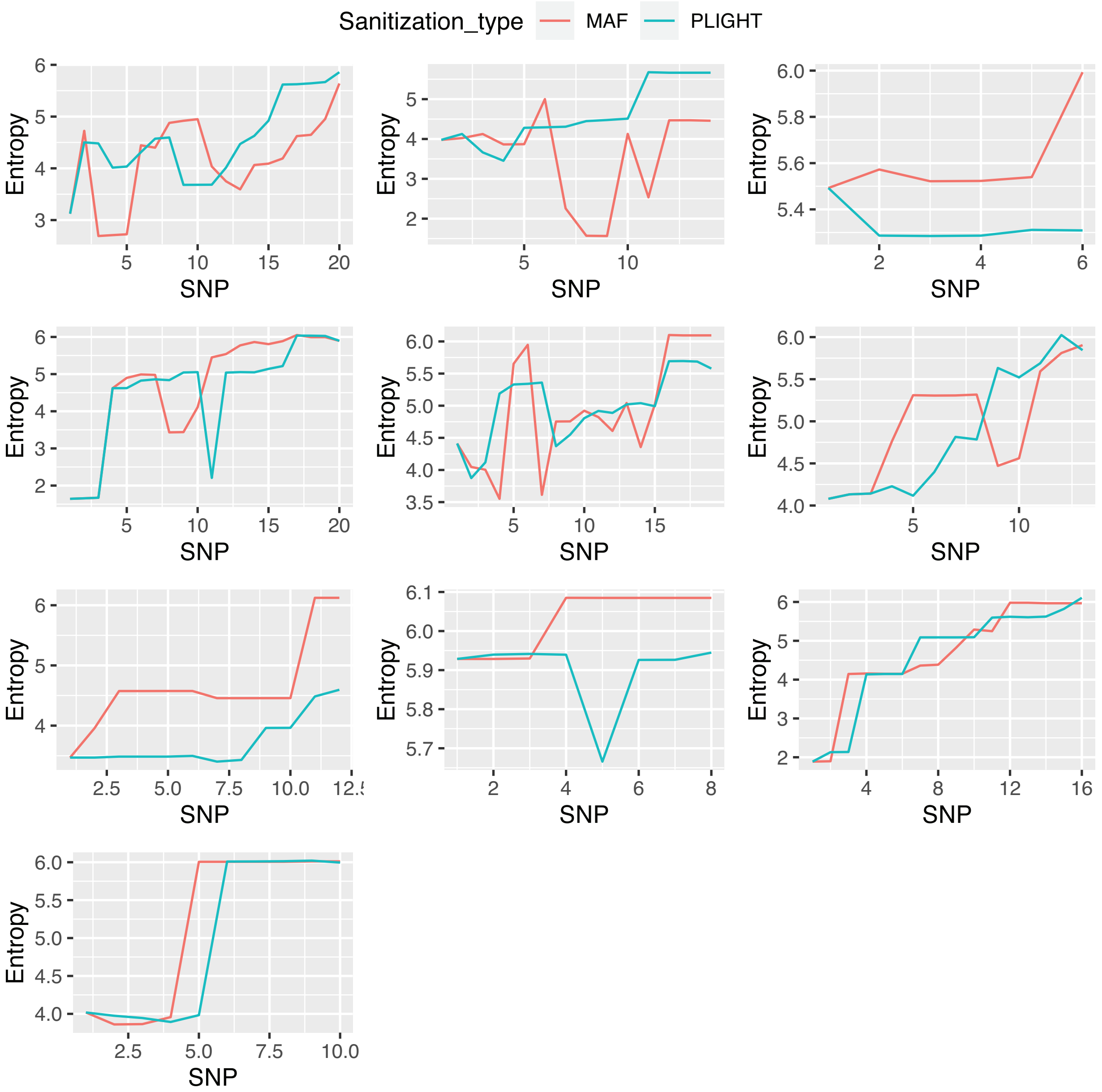


**Supplemental Figure S9.** Plot of the total entropy of individuals across all trajectories ($S_{Ind}$ defined in the main manuscript) for each of 10 independent query SNP sets, for the two different sanitization strategies. The entropy is plotted as a function of the number of SNPs removed, which varies from run to run.

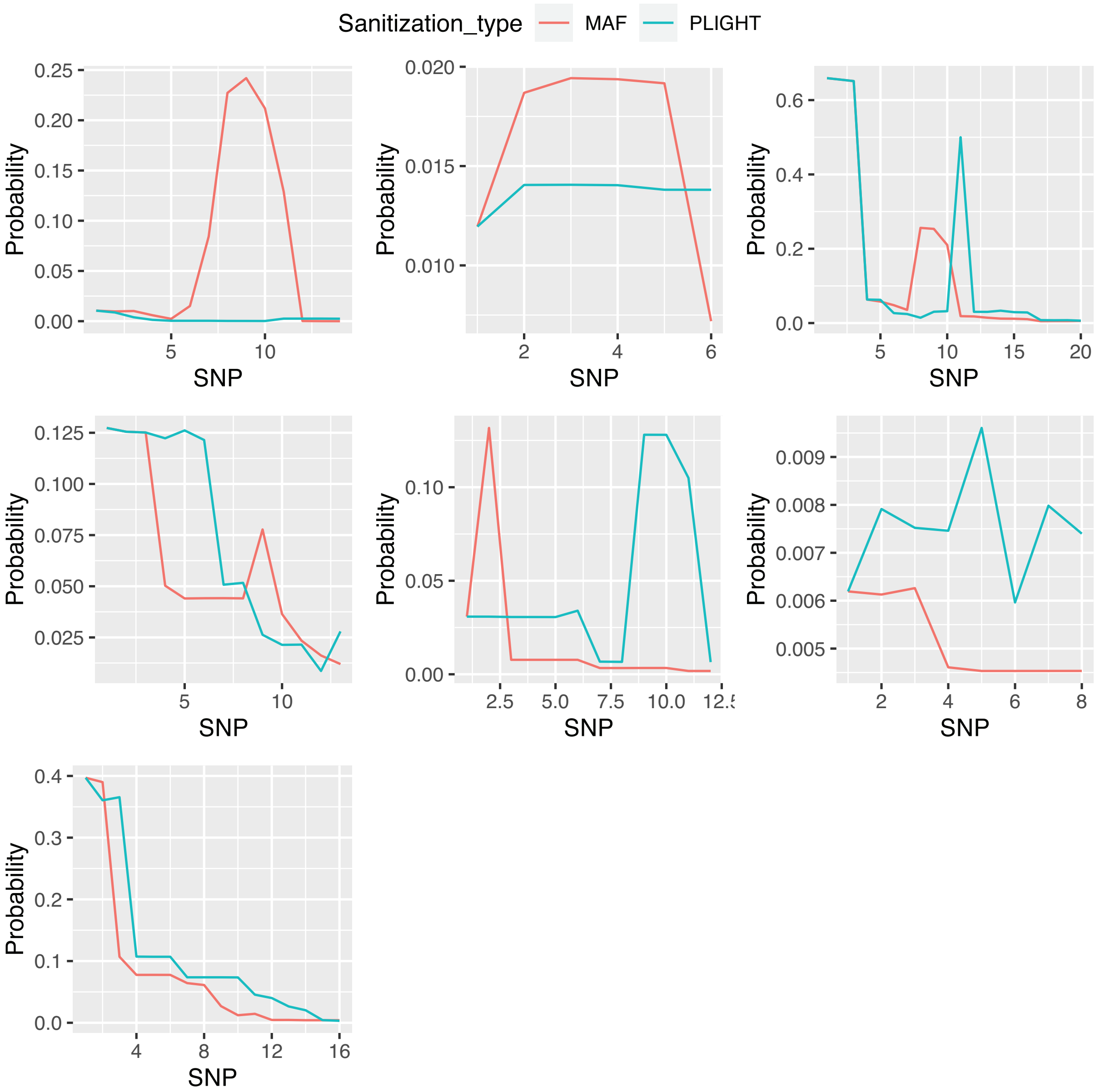


**Supplemental Figure S10.** Plot of the maximum probability of finding any of the source individuals in all trajectories ($P_{Max}^{Source}$ defined in the main manuscript) for each of 7 independent query SNP sets, for the two different sanitization strategies. 10 sets were run originally but 3 of the runs did not find the underlying source individuals. The probability is plotted as a function of the number of SNPs removed, which varies from run to run.

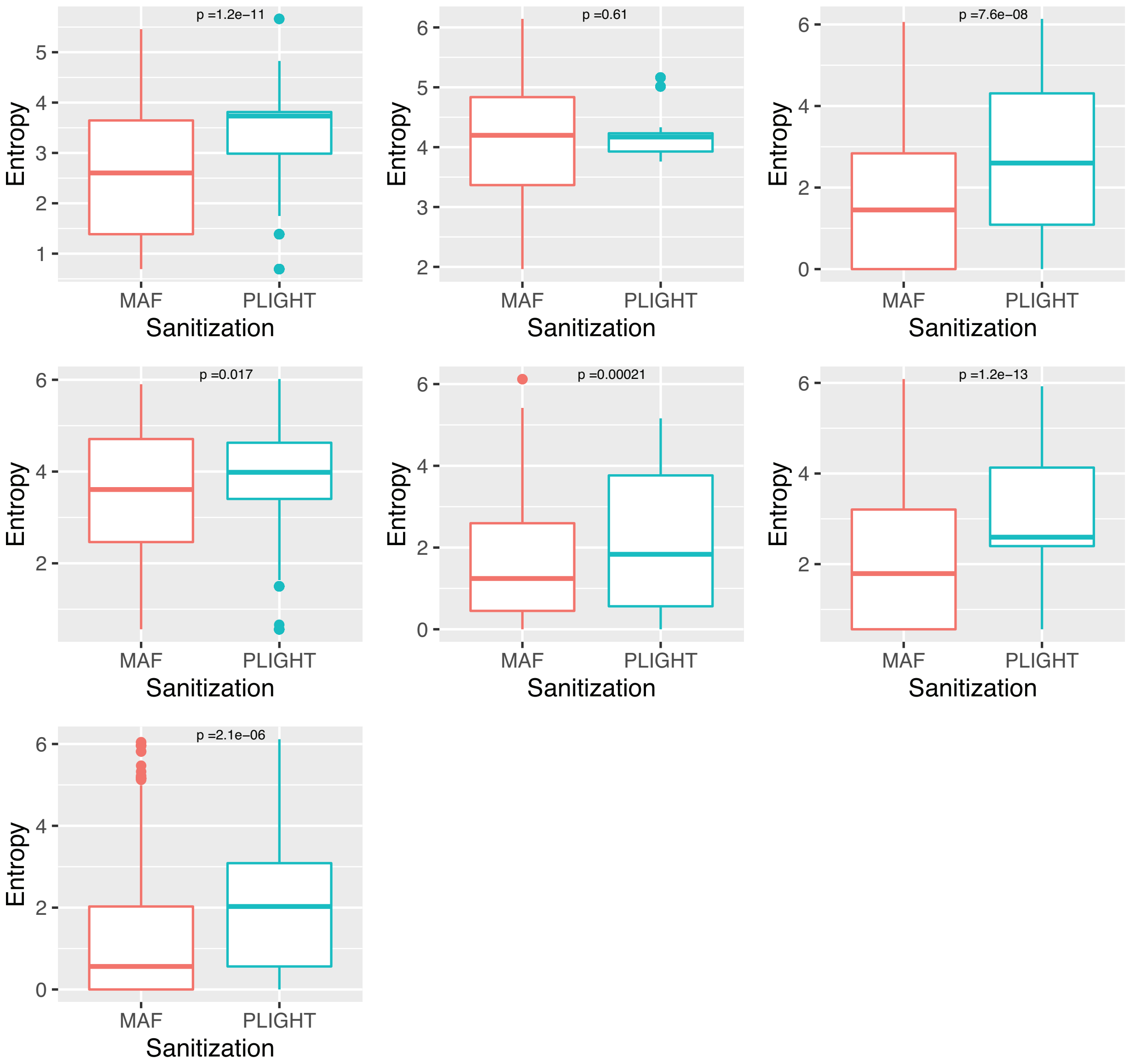


**Supplemental Figure S11.** Plot of the per-SNP entropy for those SNPs containing any of the source individuals in all trajectories ($S_{Per-SNP}(i)$ defined in the main manuscript) for each of 7 independent query SNP sets, as a function of the two different sanitization strategies. 10 sets were run originally but 3 of the runs did not find the underlying source individuals. All the per-SNP entropies are grouped together according to the sanitization strategy employed in this figure (see Supplemental Figure S12 for all the removed SNPs treated separately). P-values are shown for the comparison of means between the two distributions based on the Wilcoxon two-sample test.

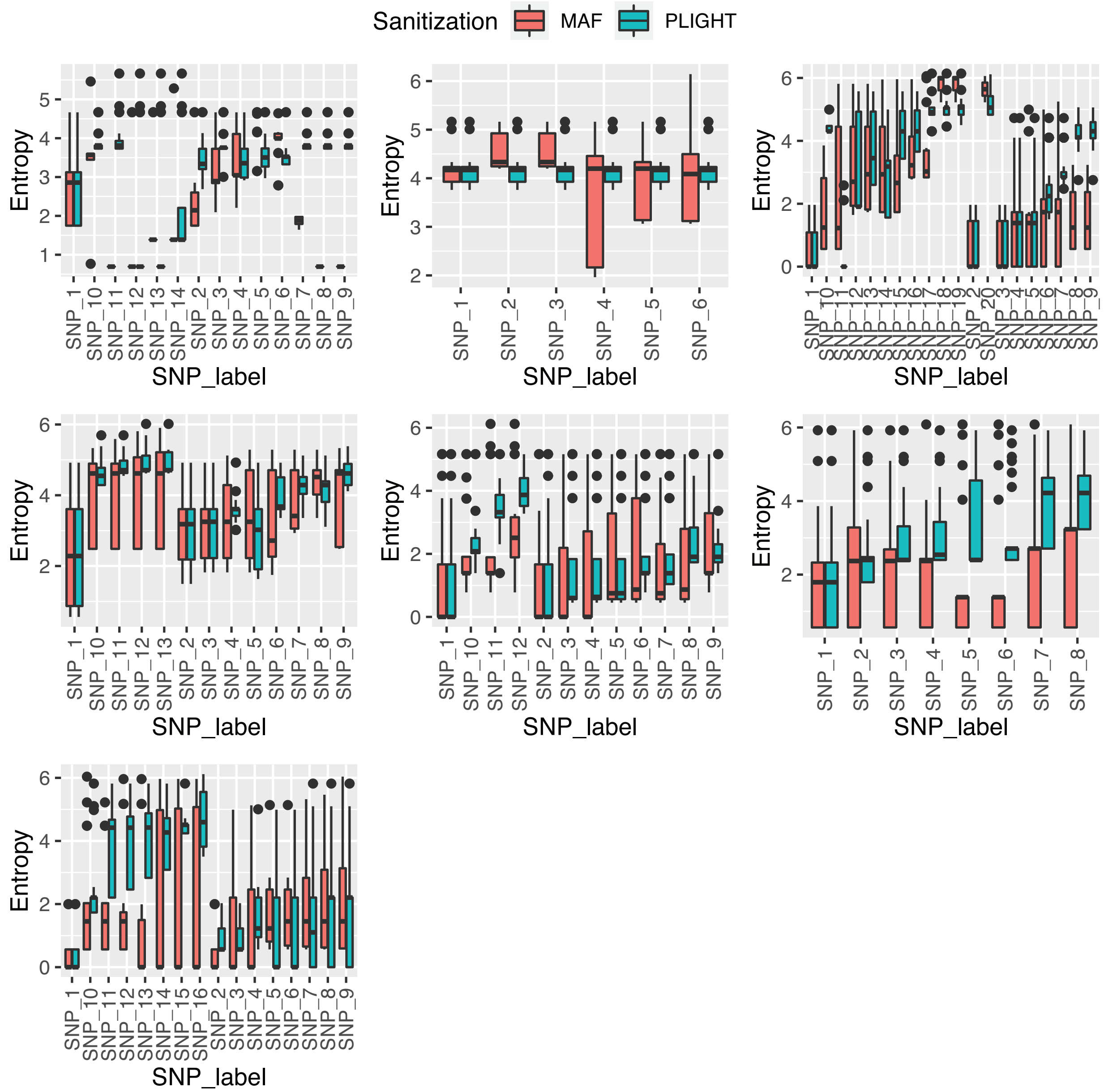


**Supplemental Figure S12.** Plot of the per-SNP entropy for those SNPs containing any of the source individuals in all trajectories ($S_{Per-SNP}(i)$ defined in the main manuscript) for each of 7 independent query SNP sets, as a function of the individual SNPs removed. 10 sets were run originally but 3 of the runs did not find the underlying source individuals.

### Supplemental Files

**Supplemental File S1.** This file reports the results of the contamination analysis, for the different mutation rates reported in Table 2 of the main manuscript and for the two populations used for the contamination simulation: (a) a "General" population where contamination was by samples randomly selected from across the 1000 Genomes cohort and (b) a "CDX" population where contamination was by samples randomly selected from the CDX population specifically. For each mutation rate, we report: (1) The minimum number of SNPs required for correct identification, in all runs (out of a maximum of 30) where the correct source individual was found; (2) The difference in the log probability of the observed SNPs between the HMM model and an independent SNP model with genotype frequencies ($\frac{log(P_{HMM})-log(P_{GF})}{N_{SNPs}}$ defined in the main text), in all runs (out of a maximum of 30) where the correct source individual was found; (3) The average minimum number of SNPs and the standard deviation (using the numbers quoted above); and (4) The average difference in the log probability and the standard deviation (using the numbers quoted above).

### Supplemental Tables

**Supplemental Table S1.** Table of conditional probabilities for the observed genotypes based on the sum of two reference haplotypes.

| | | $G_{q,l}$ | | |
|---|---|---|---|---|
| | | **0** | **1** | **2** |
| $Z^{(1)}_{j(l),l} + Z^{(2)}_{k(l),l}$ | **0** | $(1-\lambda)^2$ | $2\lambda(1-\lambda)$ | $\lambda^2$ |
| | **1** | $\lambda(1-\lambda)$ | $\lambda^2 + (1-\lambda)^2$ | $\lambda(1-\lambda)$ |
| | **2** | $\lambda^2$ | $2\lambda(1-\lambda)$ | $(1-\lambda)^2$ |

**Supplemental Table S2:** PRS matching scores for the true query genome to the best-fit mosaic trajectories for the simulated mosaic HG00360 + HG00342 SNPs across the regions of chromosomes ranging from the first observed SNP to the last observed SNP. Corresponding matching scores for the true sample to the background genomes are shown in parentheses. Four different cosine similarity scores are chosen with the aim of elucidating potentially subtle differences in the matching depending on the choice of phenotypes or method of averaging: (1) **ALL** = Cosine similarity of the true sample score relative to the mean score, for all traits; (2) **> 1 SNP** = Cosine similarity of the true sample score relative to the mean score, for all non-zero traits with more than one GWAS SNP; (3) **PRS > 2** = Cosine similarity of the true sample score relative to the mean score, for all non-zero traits where the absolute value of the Z-score of the true PRS > 2; (4) **Compare, then average** = Cosine similarity of the true sample score **relative to the score of each trajectory, subsequently averaged**, for all traits. Non-zero traits are those for which the true sample has a non-zero PRS.

| **Similarity score** | **Chromosome 1** | **Chromosome 2** | **Chromosome 21** |
|---|---|---|---|
| ***ALL:*** *Best-fit mosaics (background individuals)* | 0.9983 (0.9982) for 343 non-zero traits | 0.9694 (0.8575) for 225 non-zero traits | 0.9213 (0.9478) for 131 non-zero traits |
| ***> 1 SNP:*** *Best-fit mosaics (background individuals)* | 0.9288 (0.7637) for 197 traits | 0.9111 (0.7824) for 93 traits | 0.9517 (0.9526) for 53 traits |
| ***PRS > 2:*** *Best-fit mosaics (the background individuals were used to calculate the Z-scores)* | 0.8143 for 12 traits | 0.7683 for 14 traits | 0.9712 for 17 traits |
| ***Compare, then average:*** *Best-fit mosaics (background individuals)* | 0.9978 (0.7113) | 0.9276 (0.7811) | 0.8961 (0.8402) |

**Supplemental Table S3:** PRS matching scores for the true query genome to the best-fit mosaic trajectories for 30 and 90 coffee cup SNPs across the regions of chromosomes ranging from the first observed SNP to the last observed SNP. Corresponding matching scores for the true sample to the background genomes are shown in parentheses. Four different cosine similarity scores are chosen with the aim of elucidating potentially subtle differences in the matching depending on the choice of phenotypes or method of averaging: (1) **ALL** = Cosine similarity of the true sample score relative to the mean score, for all traits; (2) **> 1 SNP** = Cosine similarity of the true sample score relative to the mean score, for all non-zero traits with more than one GWAS SNP; (3) **PRS > 2** = Cosine similarity of the true sample score relative to the mean score, for all non-zero traits where the absolute value of the Z-score of the true PRS > 2; (4) **Compare, then average** = Cosine similarity of the true sample score **relative to the score of each trajectory, subsequently averaged**, for all traits. Non-zero traits are those for which the true sample has a non-zero PRS.

| **Cosine similarity metric** | **Chromosome 3** | **Chromosome 6** |
|---|---|---|
| ***ALL:*** *Best-fit mosaics (background individuals)* | **30-SNP-case:** 0.9975 (0.9681) for 534 non-zero traits | **30-SNP-case:** 0.8230 (0.9611) for 663 non-zero traits |
| | **90-SNP-case:** 0.9954 (0.9840) for 587 non-zero traits | **90-SNP-case:** 0.9997 (0.9766) for 690 non-zero traits |
| ***> 1 SNP:*** *Best-fit mosaics (background individuals)* | **30-SNP-case:** 0.9995 (0.9995) for 294 traits | **30-SNP-case:** -0.013 (0.9971) for 374 traits |
| | **90-SNP-case:** 0.9997 (0.9997) for 317 traits | **90-SNP-case:** 0.9992 (0.9982) for 388 traits |
| ***PRS > 2:*** *Best-fit mosaics (the background individuals were used to calculate the Z-scores)* | **30-SNP-case:** 0.5182 for 53 traits | **30-SNP-case:** -0.1138 for 78 traits |
| | **90-SNP-case:** 0.5499 for 55 traits | **90-SNP-case:** -0.0925 for 65 traits |
| ***Compare, then average:*** *Best-fit mosaics (background individuals)* | **30-SNP-case:** 0.9975 (0.8693) | **30-SNP-case:** 0.8230 (0.7494) |
| | **90-SNP-case:** 0.9881 (0.9168) | **90-SNP-case:** 0.9997 (0.7711) |

**Supplemental Table S4:** 300-SNP case. PRS matching scores for the true query genome to the best-fit mosaic trajectories for the coffee cup SNPs across the regions of chromosomes ranging from the first observed SNP to the last observed SNP. Corresponding matching scores for the true sample to the background genomes are shown in parentheses. Four different cosine similarity scores are chosen with the aim of elucidating potentially subtle differences in the matching depending on the choice of phenotypes or method of averaging: (1) **ALL** = Cosine similarity of the true sample score relative to the mean score, for all traits; (2) **> 1 SNP** = Cosine similarity of the true sample score relative to the mean score, for all non-zero traits with more than one GWAS SNP; (3) **PRS > 2** = Cosine similarity of the true sample score relative to the mean score, for all non-zero traits where the absolute value of the Z-score of the true PRS > 2; (4) **Compare, then average** = Cosine similarity of the true sample score **relative to the score of each trajectory, subsequently averaged**, for all traits. Non-zero traits are those for which the true sample has a non-zero PRS.

| Cosine similarity metric | Chromosome 3 | Chromosome 6 |
|---|---|---|
| ***ALL:*** *Best-fit mosaics (background individuals)* | **300-SNP-case:** 0.9788 (0.9840) for 596 non-zero traits | **300-SNP-case:** 0.8466 (0.9790) for 693 non-zero traits |
| ***> 1 SNP:*** *Best-fit mosaics (background individuals)* | **300-SNP-case:** 0.9995 (0.9996) for 322 traits | **300-SNP-case:** 0.8071 (0.9971) for 374 traits |
| ***PRS > 2:*** *Best-fit mosaics (the background individuals were used to calculate the Z-scores)* | **300-SNP-case:** 0.3991 for 56 traits | **300-SNP-case:** 0.0465 for 75 traits |
| ***Compare, then average:*** *Best-fit mosaics (background individuals)* | **300-SNP-case:** 0.9719 (0.8704) | **300-SNP-case:** 0.4334 (0.6846) |

### Supplemental Methods

#### Li-Stephens model and associated biological parameters

For clarity, we summarize the primary aspects of the Li-Stephens model as applicable to the work herein. Let $G_q = \{G_{q,l}\}_{l=1}^{L}$ be the genotypes of a query individual $q$, observed at SNP loci $l = \{1,2,\cdots,L\}$. The probability of observing such an individual given a space of reference haplotypes $H = \{Z_{j,l}\}_{l=1;j=1}^{l=L_{Ref};j=N}$ ($L_{Ref}$ = total number of genotyped sites in the reference genomes, $N$ = total number of haplotypes in the reference database) is:

$$P(G_q|H) = \sum_{Z_j^{(1)},Z_k^{(2)}} P\left(G_q\middle|Z_j^{(1)},Z_k^{(2)}\right).P\left(Z_j^{(1)},Z_k^{(2)}\middle|H\right) \quad \text{(S1)}$$

where the set of all possible haplotypes at the observed loci on the two chromosomes is given by $Z_j^{(\alpha)} = \left\{Z_{j(l),l}^{(\alpha)}\right\}_{l=1}^{L}$, with $Z_{j(l),l}^{(\alpha=1,2)}$ being the haplotype at position $l$, and $j$ being the index of the sampled haplotype. We treat the haplotype index $j(l)$ as a function of $l$, as it is possible for the choice of reference haplotype to be different at each locus; that is, in the haplotype matching process, recombination between reference haplotypes may occur from one observed locus to the next (see **Figure S1**). The second subscript explicitly indicates that, for reference haplotype $j(l)$, we select the genotype at locus $l$.

The assumption in the current iteration of the algorithm is that the observed genotypes and the reference haplotypes are registered with respect to the same, linear reference genome. This enables a simpler matching of reference haplotypes to observed genotypes. For data structures such as personal genomes and graph genomes, additional genotype matching strategies would need to be incorporated, but the conceptual framework of searching through recombining haplotypes would be the same. In general, the set of genotyped sites does not have to perfectly overlap with the set of reference haplotype sites because of rare SNPs in an individual's genotype or differences in genotyping arrays. However, for the purposes of this study we only consider genotyped sites that overlap with those of the reference haplotypes, especially given our interest in determining the identification power of common SNPs. In case of the presence of structural variants overlapping SNP loci, we allow for missing loci in any of the reference haplotypes. We thus consider the reference haplotypes as providing the complete search space, especially in light of the constantly growing genetic databases available for comparison. We avoid making explicit assumptions of population membership and statistics for the query individual with the belief that, beyond the implicit assumptions of the chosen reference set, this will enable more unbiased estimates of kinship and genotypic similarity.

$P\left(Z_j^{(1)},Z_k^{(2)}\middle|H\right)$ contains information on the "trajectories" through haplotype space that emerge from the reference set: it is the probability of obtaining a given set of haplotype observations at all the query loci. In general, this is not a simple measure of the frequencies of entire reference haplotypes (unless the query genotype is known to be in the reference set) due to the possibility of recombination. Recombination is incorporated into the analysis in the expressions for the transition probabilities from one query site to the next. Using results of Li and Stephens (Li and Stephens 2003) and Marchini et al (Marchini et al. 2007), we have

$$P\left(Z_j^{(1)}, Z_k^{(2)} \middle| H\right) = P\left(Z_{j(1),1}^{(1)}, Z_{k(1),1}^{(2)} \middle| H\right) \prod_{l=1}^{L-1} P\left(\left\{Z_{j(l),l}^{(1)}, Z_{k(l),l}^{(2)}\right\} \rightarrow \left\{Z_{j(l+1),l+1}^{(1)}, Z_{k(l+1),l+1}^{(2)}\right\} \middle| H\right) \qquad \text{(S2)}$$

where $P\left(Z_{j(1),1}^{(1)}, Z_{k(1),1}^{(2)} \middle| H\right)$ is the probability of observing a given set of haplotypes at the first query locus. This is often drawn from a uniform distribution across all haplotype pairs, but could be modified if prior knowledge on the membership of the query individual in a particular subpopulation is available. The transition from one site to the next is given by

$$P\left(\left\{Z_{j(l),l}^{(1)}, Z_{k(l),l}^{(2)}\right\} \rightarrow \left\{Z_{j(l+1),l+1}^{(1)}, Z_{k(l+1),l+1}^{(2)}\right\} \middle| H\right) =$$

$$\begin{cases} \left(e^{-\frac{\rho_l}{N}} + \frac{1-e^{-\frac{\rho_l}{N}}}{N}\right)^2 & \mathit{if\ neither\ haplotype\ changes\ from\ site\ } l \mathit{\ to\ } l+1 \\ \left(e^{-\frac{\rho_l}{N}} + \frac{1-e^{-\frac{\rho_l}{N}}}{N}\right)\left(\frac{1-e^{-\frac{\rho_l}{N}}}{N}\right) & \mathit{if\ one\ haplotype\ changes, but\ not\ the\ other} \\ \left(\frac{1-e^{-\frac{\rho_l}{N}}}{N}\right)^2 & \mathit{if\ both\ haplotypes\ change} \end{cases} \qquad \text{(S3)}$$

with $\rho_l = 4N_e r_l$, $r_l$ being the per generation genetic distance between the sites $l$ and $l+1$. The exponential dependence arises from the assumption of a Poisson process of recombination at any position in the genome, while the division by $N$ in the exponent ensures that the probability of a recombination event drops exponentially with growth in the reference haplotype database. The latter idea ensures that the number of truly novel haplotypes reaches a plateau, i.e. there is a stable set of possible haplotypes in a population. In terms of the recombination rate $c_l$ and the physical distance between adjacent loci $d_{l\rightarrow l+1}$, $r_l = c_l \times d_{l\rightarrow l+1}$. $N$ is the number of reference haplotypes, and $N_e$ is the effective population size, taken to be 11,418 (Li and Stephens 2003; Marchini et al. 2007; International HapMap Consortium 2003).

In our methods, the linear model $\rho_l = 4N_e c_l d_{l\rightarrow l+1}$ is included as the default. However, we allow for the inclusion of a user-defined model of recombination in its place. For example, if there is a known recombination hotspot between two adjacent query sites, it would alter the probability of transitioning between reference haplotypes in the search space, and thus impact the best-fit haplotypes calculated by the method. The user can explicitly include a vector of recombination values to be used for the $L-1$ intervals between query sites.

Another essential aspect of Equation S3 is the implicit assumption of uniform transition probabilities. That is, in its current form, Equation S3 does not discriminate between transitions from one haplotype to any other. If, on the other hand, a model is to be constructed where different subgroups have distinct recombination rates at particular locations and/or are assumed to impacted by assortative mating then transition probabilities would be conditional based on membership in these subgroups. In this iteration of our model, we do not provide a framework of this nature, but such an update would simply require the inclusion of appropriate bias terms conditional on the memberships of the initial and final haplotypes. However, we wish to emphasize that maintaining uniform transition probabilities helps prevent biased interpretations of ethnic group membership and isolation, and allows for the broad intermixing of haplotypes known to have occurred throughout human history(Narasimhan et al. 2019).

The other term in Equation S1, $P\left(G_q \middle| Z_j^{(1)}, Z_k^{(2)}\right)$, quantifies the probability of observing the query genotypes given a particular set of underlying haplotypes. This probability helps constrain the haplotypes that are possible given the observed genotypes, allowing for the case where mutations or genotyping errors occur (as considered in IMPUTE(Howie et al. 2009)).

$$P\left(G_q \middle| Z_j^{(1)}, Z_k^{(2)}\right) = \prod_{l=1}^{L} P\left(G_{q,l} \middle| Z_{j(l),l}^{(1)}, Z_{k(l),l}^{(2)}\right) = \prod_{l=1}^{L} P\left(Z_{j(l),l}^{(1)} + Z_{k(l),l}^{(2)} \to G_{q,l}\right) \quad \text{(S4)}$$

with $P\left(Z_{j(l),l}^{(1)} + Z_{k(l),l}^{(2)} \to G_{q,l}\right)$ determined by the number of sites that require mutation to match the observed genotypes shown in **Table S1**.

We follow the suggestion of the authors of IMPUTE to consider a background rate of base pair mutation $\theta$ that translates into a mutation rate per haplotype of $\lambda = \frac{\theta}{2(N+\theta)}$ under the assumption of a neutral coalescent tree for $N$ haplotypes(Li and Stephens 2003; Marchini et al. 2007). However, it is possible for the user to explicitly augment the background rate $\theta$ with contributions from genotyping error, or to ignore the mutation rate altogether and set $\theta = \frac{2N\lambda}{1-2\lambda}$ such that $\lambda$ is equal to the known genotyping error. Thus, our code allows for either $\theta$ (the ***thetamutationrate*** parameter) or $\lambda$ (the ***lambdamutationrate*** parameter) to be set.

To summarize, the aim is to figure out the contribution to the total probability of each of the haplotype combinations, by estimating $P\left(G_q \middle| Z_j^{(1)}, Z_k^{(2)}\right).P\left(Z_j^{(1)}, Z_k^{(2)} \middle| H\right)$ for all haplotype trajectories, and to maximize this probability.

#### Hidden Markov Model optimization

The problem of identifying the best-fit combination of haplotypes is well-suited to the framework of Hidden Markov models (HMMs) given the traditional treatment of the genome as a linear sequence of base pairs. In this understanding, meiotic recombination between loci does not occur between distant locations of a chromosome (as may occur, hypothetically, due to consistent 3D folding of the chromosomes within the nucleus), but has a certain probability of occurring at every intermediate site between any pair of loci. Usually, the greater the distance between the loci, the higher is the probability that recombination will have occurred in an ancestor of the query genome, though the probability is not necessarily uniform across every site. As seen in the previous section, it then becomes easy to associate HMM emission probabilities at genomic sites with mutation rates and HMM transition probabilities between latent haplotypes with recombination rates. Furthermore, in the above expressions first-order Markovian behavior is assumed, and the observed output genotype is seen to depend only on the underlying haplotypes at that site alone (so-called output independence). This constrains the type of HMMs considered here, but leaves open interesting future applications where such assumptions are relaxed.

Accordingly, the problem of identifying the best trajectory through haplotype space can be carried out using the Viterbi algorithm (Viterbi 1967). This method solves the problem of maximizing the probability of the trajectories through the latent space in time $O((N \times N)^2 L)$, where $N$ is the number of possible haploid states, i.e. the number of reference haplotypes, $N \times N$ is the corresponding number of diploid states, and $L$ is the number of observed loci:

$$Most\ likely\ trajectory = \underset{Z_j^{(1)}, Z_k^{(2)} \in \left\{Z_{j(l),l}^{(1,2)}\right\}_{l=1;j=1}^{l=L;j=N}}{\operatorname{argmax}} P\left(G_q \middle| Z_j^{(1)}, Z_k^{(2)}\right).P\left(Z_j^{(1)}, Z_k^{(2)} \middle| H\right)$$

$$= \underset{Z_j^{(1)}, Z_k^{(2)} \in \left\{Z_{j(l),l}^{(1,2)}\right\}_{l=1;j=1}^{l=L;j=N}}{\operatorname{argmax}} P\left(G_q, Z_j^{(1)}, Z_k^{(2)} \middle| H\right)$$

(S5)

where expressions for the two probabilities are given in Equations S2, S3 and S4.

The Viterbi algorithm achieves a more efficient solution to the optimization problem in equation S5 than the naïve search by recognizing that the overall optimization problem can be separated into separate optimization steps at each query site with the convenient iterative scheme:

$$P\left(\left\{G_{q,\delta}\right\}_{\delta=1}^{l}, \left\{Z_{j(\delta),\delta}^{(1)}, Z_{k(\delta),\delta}^{(2)}\right\}_{\delta=1}^{l-1}, Z_{j(l),l}^{(1)}, Z_{k(l),l}^{(2)} \middle| H\right) =$$

$$P\left(G_q \middle| Z_{j(l),l}^{(1)}, Z_{k(l),l}^{(2)}, H\right). \max_{j(l-1),k(l-1)} \begin{bmatrix} P\left(\left\{G_{q,\delta}\right\}_{\delta=1}^{l}, \left\{Z_{j(\delta),\delta}^{(1)}, Z_{k(\delta),\delta}^{(2)}\right\}_{\delta=1}^{l-2}, Z_{j(l),l}^{(1)}, Z_{k(l),l}^{(2)} \middle| H\right) \times \\ P\left(Z_{j(l-1),l-1}^{(1)} \to Z_{j(l),l}^{(1)}, Z_{k(l-1),l-1}^{(2)} \to Z_{k(l),l}^{(2)} \middle| H\right) \end{bmatrix}$$

$$\Rightarrow v_l\big(j(l), k(l)\big) = EmissionProb\left(G_{q,l} \middle| Z_{j(l),l}^{(1)}, Z_{k(l),l}^{(2)}, H\right) \times$$

$$\max_{j(l-1),k(l-1)} \left[v_{l-1}\big(j(l-1), k(l-1)\big).TransitionProb\left(Z_{j(l-1),l-1}^{(1)} \to Z_{j(l),l}^{(1)}, Z_{k(l-1),l-1}^{(2)} \to Z_{k(l),l}^{(2)} \middle| H\right)\right]$$

with $v_l\big(j(l), k(l)\big) = P\left(\left\{G_{q,\delta}\right\}_{\delta=1}^{l}, \left\{Z_{j(\delta),\delta}^{(1)}, Z_{k(\delta),\delta}^{(2)}\right\}_{\delta=1}^{l-1}, Z_{j(l),l}^{(1)}, Z_{k(l),l}^{(2)} \middle| H\right)$

(S6)

where the algorithm initializes a value of $v_1\big(j(1), k(1)\big)$ and then proceeds to iteratively update the probability. The second line of Equation S6 simply provides a clearer conceptual understanding of the first line.

$P\left(\left\{G_{i,\delta}\right\}_{\delta=1}^{l}, \left\{Z_{j(\delta),\delta}^{(1)}, Z_{k(\delta),\delta}^{(2)}\right\}_{\delta=1}^{l-1}, Z_{j(l),l}^{(1)}, Z_{k(l),l}^{(2)} \middle| H\right)$ is the joint probability of the observed genotypes at all sites up to and including site $l$, and of the reference haplotypes at all sites up to site $l-1$, with the reference haplotypes at site $l$ being fixed at $Z_{j(l),l}^{(1)}, Z_{k(l),l}^{(2)}$, and is easily related to the argument of the *argmax* function on the right-hand side of equation S5.

***Truncation scheme.*** In the *PLIGHT_Truncated* module, at every observed site we truncate the possible haplotype states by choosing the top $T$ sets of $(\alpha, \beta)$ pairs. This scheme was inspired by similar techniques employed in the Eagle2 imputation program (Loh et al. 2016). The main premise of this approximation is after a certain point, only a fraction of a the total number of trajectories will meaningfully contribute to the best-fit states, and allow the retention of only a fraction of the total number of states in memory. However, in addition to tracking the probabilities and backtrace vectors of these states, we now also need to retain positional indices associated with the particular pairs for future look-up (i.e. we need to know the identity of these pairs).

The module allows the user to set the truncation factor as the fraction of the total number of reference haplotype pairs retained in the calculation:

Truncation factor $f = T/\left(\frac{N(N+1)}{2}\right) = T/T_{tot}\,;\ T_{tot} = \frac{N(N+1)}{2}$

(S7)

However, we recognize that immediate truncation at the first location would not allow the probabilities of the most likely trajectories to build up sufficiently. Therefore, we phase-in the truncation by using a linear decrease in the value of $T$ until the halfway point:

$$T_l = \begin{cases} floor(slope \times (l-1) + T_{tot}), for\ l \leq floor\left(\frac{L}{2}\right) \\ floor(f \times T_{tot}), for\ l > floor\left(\frac{L}{2}\right) \end{cases}$$

(S8)

where $slope = T_{tot} \times (f-1)/\left(floor\left(\frac{L}{2}\right) - 1\right)$ and $floor(x)$ is the rounding down function. Furthermore, setting a cutoff on the number of states could possibly arbitrarily remove states that have the same probability as the included ones. Accordingly, we soften the truncation by including all additional states that have the exact same probability as the final $T_l$ state. Note that this may grow the size of the matrices significantly, if the number of equiprobability or degenerate states is large. This may occur for very sparse data.

The advantage of the truncation scheme is the reduction of the size of the matrices. To maintain this advantage, we had to explicitly calculate the probabilities at the first observed site, where the truncation had not been applied yet. This prevents (in the current version of the code) the inclusion of missing genotypes. Additionally, the parallelization procedure at the first observed site required the chunking up of the haplotype pairs into subgroups the size of $floor(f \times T_{tot})$, and running each of these subgroups through parallel matrix calculations. This prevented the necessity of having the entire set of haplotype pairs be manipulated in matrices simultaneously, which would have defeated the purpose of the truncation process.

### Supplemental Results

**Supplement to Section *"Truncated algorithm".*** We ran the same SNP set as in Section "***Exact search within a reference database of 400 haplotypes"*** through the truncated algorithm to assess the degree of compressibility of the trajectories, with a recombination rate of 0.5 cM/Mb and with a search through 200 reference individuals. In interpreting the results, it is worth bearing in mind the fact that while the algorithm was set up to slowly phase-in the restriction of the number of considered haplotype pairs to the top few, it yet retained a degree of elasticity to prevent an abrupt cut-off: all haplotype pairs with the same probability as the last state in the imposed cut-off were also included (**Supplemental Figure S4**). We ran the calculation for two different asymptotic truncation levels, with the truncation factor $f$ in Equations S7 and S8 = 0.005 and 0.02 (**Figures S4A** and **S4B**). For reference, the fractions of the matrix sizes at each SNP in the calculations are shown for $f$ = 0.005 (**Figure S4A**) and $f$ = 0.02 (**Figure S4B**). The resultant trajectories for chromosome 1 were identical to the exact algorithm, implying that reducing the number of considered trajectories did not impact the search process. For chromosome 21, the lower truncation level of 0.005 resulted in fewer trajectories being included, while the higher truncation of 0.02 reproduced the same trajectories as the exact algorithm. Chromosome 2 appeared to be less resilient to truncation and produced different results from the exact algorithm (the trajectories for $f$ = 0.005 are shown in **Figure S4C**; compare to **Figure 3B**). Note that truncation at later SNPs favors the prioritization of HG00360 in the earlier parts of the chromosome, whereas the exact algorithm with the same recombination rate removes this individual from the final results. The degree to which different trajectories are resilient to truncation is a measure of the degree to which the best trajectories separate out from the others. In informatic terms, if the best trajectory probabilities (analogous to the energies of physical states) have low entropy (analogous to deep, sharp valleys in the energy landscape) trajectory truncation will not impact the search; if the entropy is high (broad, shallow valleys in the energy landscape) truncation will impact the results.

**Supplement to Section *"Inference based on SNPs obtained from coffee cups"***

Similar to the analyses in the main manuscript, we carried out a PRS and genotype comparison for the case where 300 query SNPs are drawn from coffee cups. We first created a list of ~500 SNPs that included the 90 from the main manuscript analysis, but then sub-selected this to a more tractable set of 300 SNPs. Thus, the overlap in the 30/90 and 300 SNP sets are not perfect. However, this analysis serves as an approximate extrapolation to larger SNP set sizes. We then ran *PLIGHT_Iterative* with parameters $S_{sg} = 300, \text{mutation rate } \lambda = 0.1, \text{number of iterations } \mathrm{n}_{\mathrm{iter}} = 1.$

With a scaling up of the number of SNPs, we also noticed a concurrent increase in the total number of best-fit trajectories. We chose a subset of 10 trajectories per chromosome for the following analyses. We first evaluated the degree of genotype matching in the mosaics relative to a background population of 98 individuals. For chromosome 3, the exact-matching-genotype fractions across the 10 mosaic trajectories were in the range 0.45-0.47, compared to a background of 0.36-0.46, with a Welch's two-sample t-test p-value = 2.2e-16. The correspondence scores were $C$ = 0.32-0.34 for the mosaic trajectories, against a background of

0.29-0.33, with a p-value = 1.4e-15. For chromosome 6, the exact-matching-genotype fractions across the 10 mosaic trajectories were in the range 0.43-0.44, compared to a background of 0.34-0.43, with a Welch's two-sample t-test p-value = 2.2e-16. The correspondence scores were $C$ = 0.33-0.34 for the mosaic trajectories, against a background of 0.29-0.33, with a p-value = 2.2e-16.

We also carried out a PRS analysis for the resulting mosaics. There was no significant improvement in the inference of PRS scores, relative to the 30- and 90-SNP cases. The results are shown in Table S2.
